# methylTFR: Computational quantification of transcription factor activity from DNA methylation

**DOI:** 10.64898/2026.09.29.755279

**Authors:** Irem B. Gündüz, Regina Nitsch, Sarath Kumar Murugan, Fabian Müller

**Affiliations:** Integrative Cellular Biology & Bioinformatics Lab, Saarland Informatics Campus, Saarland University, 66123 Saarbrücken, Germany

**Keywords:** DNA methylation, Transcription factors (TFs), R package, Gene regulation

## Abstract

DNA methylation modulates transcription factor (TF) binding, and plays an important role in regulating gene expression. We developed methylTFR, an R package for quantifying TF activity from DNA methylation across TF-binding sites, providing a low-dimensional and interpretable activity profile. Across 147 human immune cell methylomes, methylTFR identified cell-type specific regulators, including CEBP and ETS family activity in myeloid cells. In CD4+ T cells, TF activity scores resolved the naive-to-memory trajectory and highlighted AP-1 factors as important memory regulators. We applied methylTFR to sparse single-cell methylome data, and we integrated derived activity scores with gene expression and accessibility into joint factor models yielding multimodal views of lineage regulators.

## Background

Transcription factors (TFs) are DNA-binding proteins that regulate transcription by recognizing target DNA sequences or motifs with varying degrees of specificity and by recruiting RNA polymerase to initiate gene expression, by recruiting cofactors and chromatin-modifying complexes that render the surrounding region permissive or repressive, or by blocking the binding region of other activators and the polymerase, thereby reducing gene expression [1]. TF-binding motifs, curated and experimentally validated, are available in large catalogs such as JASPAR2020 and CISbp-v2 and have been clustered into similar motifs [2,3,4]. Epigenetic dysregulation of transcription factor binding sites (TFBSs) is associated with abnormal gene expression, which has been implicated in crucial regulatory roles in various diseases, including viral infections [5], cancer [6], and Alzheimer’s disease [7].

Understanding the dynamics of TF activity through its interactions with DNA elements can help identify potential therapeutic targets and enhance our understanding of gene regulatory mechanisms [8]. TF activity can be computationally inferred from the genome-wide aggregation of chromatin accessibility signals across TF-binding motifs, thereby revealing the footprint of characteristic binding profiles [9,10]. However, this approach is limited to analyzing chromatin accessibility profiles around TFBSs. Yet, gene regulation is a complex mechanism orchestrated by the interplay between various epigenetic modifications and chromatin accessibility.

One such epigenetic modification is DNA methylation, in which cytosine bases are methylated, forming 5-methylcytosine (5mC). Genome-wide DNA methylation patterns can be analyzed using bisulfite or enzymatic conversion, followed by sequencing [11]. Investigating genome-wide DNA methylation patterns has enhanced our understanding of cell states and developmental trajectories, including embryonic development and human T-cell memory (Tmem) formation [12,13]. Despite advances in library preparation and sequencing protocols that now enable DNA methylation profiling of low-input samples or single cells, the resulting data often have limited coverage and density. Thus, cell-type-level investigation of DNA methylation patterns at individual loci remains challenging from these data [14].

DNA methylation plays a crucial role in regulating cell states and influences gene expression by modulating TF binding to their specific binding sites (TFBS) [15]. Typically, transcriptional repression is associated with CpG dinucleotide methylation in gene promoters. While many studies focus on genes or promoters as units for comparing cell states, most of the lineage-associated variance is explained by DNA methylation in gene-distal regions [16]. The role of DNA methylation in regulating transcription factor binding sites in these distal regulatory elements, such as enhancers, can be more complex [16]. DNA methylation in these regions can exert both positive and negative effects on transcription factor binding affinity [16]. Hence, comprehensively characterizing DNA methylation patterns at transcription factor binding sites and their interplay with chromatin accessibility is necessary to improve understanding of TF binding dynamics.

An example of epigenetic plasticity at TFBS and its potential to regulate cell state is given by T cell memory formation. T helper (Th) cells play a crucial role in the adaptive immune response, providing long-term protection against pathogens [17]. They exist in different memory cell states, such as central memory (Tcm), effector memory (Tem), and terminally differentiated (Temra) cells. Changes in DNA methylation at transcriptional elements can be used to identify potential lineage regulators and to model human CD4+ T memory cell differentiation [13]. Furthermore, Durek et al. (2016) reported segmental loss of DNA methylation within global methylation patterns during Tmem cell differentiation.

To analyze cell state through the lens of DNA methylation profiles in candidate regulatory elements (cCREs), we developed methylTFR, an R package that quantifies transcription factor activity from DNA methylation signatures at transcription factor binding sites, derived from bulk or aggregated single-cell experiments, yielding insights into the genome-wide regulatory implications of TFs. Quantifying TF activity this way also facilitates integrating DNA methylation with other regulatory layers, such as chromatin accessibility and gene expression, using factor models. We demonstrate the utility of methylTFR across three different use cases. First, in 147 WGBS samples covering 14 human immune cell types, 632 motif activity scores classified cell type at least as accurately as 2.7 million 1-kb tiling regions, and the CEBP and ETS families separated the myeloid from the lymphoid compartment. Second, in CD4+ T cells, the deviation scores ordered naive, central memory, and effector memory populations along the trajectory of memory formation and revealed a shift from SPI and ETS activity toward AP-1 that agreed with motif enrichment in differentially methylated regions in the same comparison. Third, in pseudobulk profiles from a single-cell multimodal dataset of immune cells under viral and chemical exposure, methylation- and chromatin-accessibility-derived activities were negatively correlated for most factors, and methylTFR resolved differences between T-cell states that were not apparent from accessibility alone.

## Results

### methylTFR – an R package for quantifying transcription factor (TF) motif activity from DNA methylation profiles

Transcription factor activity has previously been defined from gene expression, co-expressed gene modules [18], or from chromatin accessibility by aggregating ATAC-seq or DNase-seq signals across the binding motifs of a given factor [9,10]. This confines the analysis to a single regulatory layer and ignores DNA methylation, which has been shown to modulate TF binding affinity at binding sites [11,16]. methylTFR addresses this gap by inferring TF activity directly from DNA methylation, providing a standardized, quantitative way to summarize DNA methylation at binding motifs and to report methylation-derived activity per TF rather than per locus, thereby keeping the output interpretable, relatively low-dimensional through genome-wide aggregation, and directly comparable to accessibility- and expression-based measures (Figure 1).

**Figure 1:**
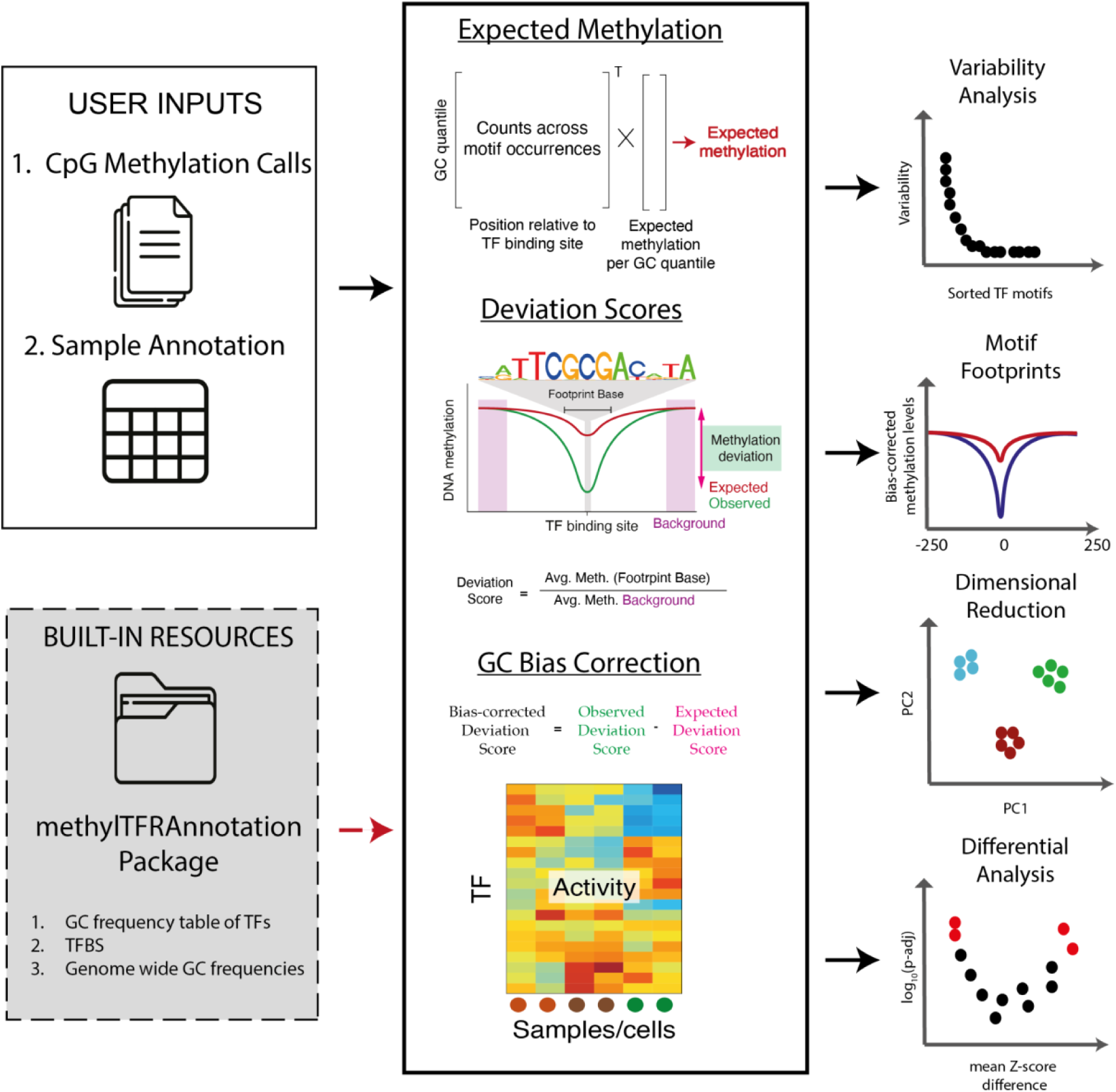
Overview of the methylTFR framework. methylTFR quantifies transcription factor activity by aggregating DNA methylation across TFBS. The inputs are CpG methylation calls from bulk or aggregated single-cell experiments, corresponding sample annotations, and TBFS annotations from the methylTFRAnnotation package. From these, methylTFR computes per-motif footprint scores and corrects for GC content to derive bias-corrected deviation scores that quantify motif-specific TF activity across samples and are suitable for downstream analysis.

We implemented methylTFR in the R framework compatible with other packages in the Bioconductor [19] environment (Figure 1). methylTFR requires (i) CpG-level DNA methylation calls from bulk or single-cell experiments and (ii) a sample annotation file. From these inputs, methylTFR computes DNA methylation deviation scores, corrects for GC bias, and derives bias-corrected scores that capture motif-specific TF activity across samples (see Methods). Genome-assembly-specific annotations are stored in the auxiliary methylTFRAnnotation packages. These packages are currently available for genome assemblies GRCh38 (Hg38) and Mus musculus (mm10), and they contain: (i) motif-specific GC content frequency table (GCfreqs), (ii) transcription factor binding site (TFBS) regions derived from reference motif sets (JASPAR2020, Altius, CISbp-v2) [2,3,4], and (iii) genome-wide GC distribution table (gcDist) discretized into five frequency quantile levels. The methylTFR algorithm uses these annotations to summarize methylation counts across TFBS from observed methylation levels in a given sample and to compute expected methylation levels based on motif GC content (see Methods). For flexibility, we also provide methylTFRAnnotationBuilder, an R package that enables users to generate custom annotation files for any genome assembly of interest (see Supplementary packages).

The motif-specific TF activity scores can then be used for exploratory and comparative analyses, all of which are implemented in the package, including dimensionality reduction (e.g., PCA), differential activity testing between phenotypes or conditions, variability analysis across TF motifs, and motif-specific footprint visualization.

### methylTFR quantifies TF activity across human immune cells and captures cell-type-dependent regulatory signatures

We applied methylTFR to whole-genome bisulfite sequencing (WGBS) data from 147 samples representing 14 distinct human immune cell types, including NK cells, monocytes, B cells, plasma cells, and T cells (BLUEPRINT project, Figure 2A) [20]. TF activity was quantified across 632 motifs annotated in the JASPAR2020 database [3]. We compared the low-dimensional embeddings of methylTFR deviation scores to those derived from average DNA methylation in tiling regions across 1-kb genomic windows (2,674,318 regions) or in regions annotated as distal enhancers (74,554 regions) (Figure 2B) [20]. Principal components derived from 632 motif activity scores separated immune cell types more effectively than those from tiling regions, which failed to distinguish several myeloid and lymphoid populations in the first two principal components (Figure 2B). This separation depended on the bias correction: with raw, uncorrected deviation scores, cell types overlapped in the first two components, and the embedding explained less variance (Supplementary Figure 1A). The correction also interacts with which regions are scored; restricting the count to distal sites resolved the populations better than scoring every TFBS in the genome-wide JASPAR set (Figure 2B, Supplementary Figure 1A). The two approaches also differ in how their variance is spread: for the tiling and distal methylation, the first component alone carried 99.0% and 98% of the variance, whereas for methylTFR it was shared across the first two components, resulting in an embedding with more than one informative axis (Supplementary Figure 1B). Promoters and gene bodies exhibit methylation levels that are highly consistent across cell types, while distal regulatory elements are more variable. Motif matches that fall within promoters therefore add a signal that barely changes between samples, which likely dilutes the cell-type differences carried by the distal sites [17,22]. All further analyses therefore use bias-corrected deviation scores computed over distal regions.

**Figure 2:**
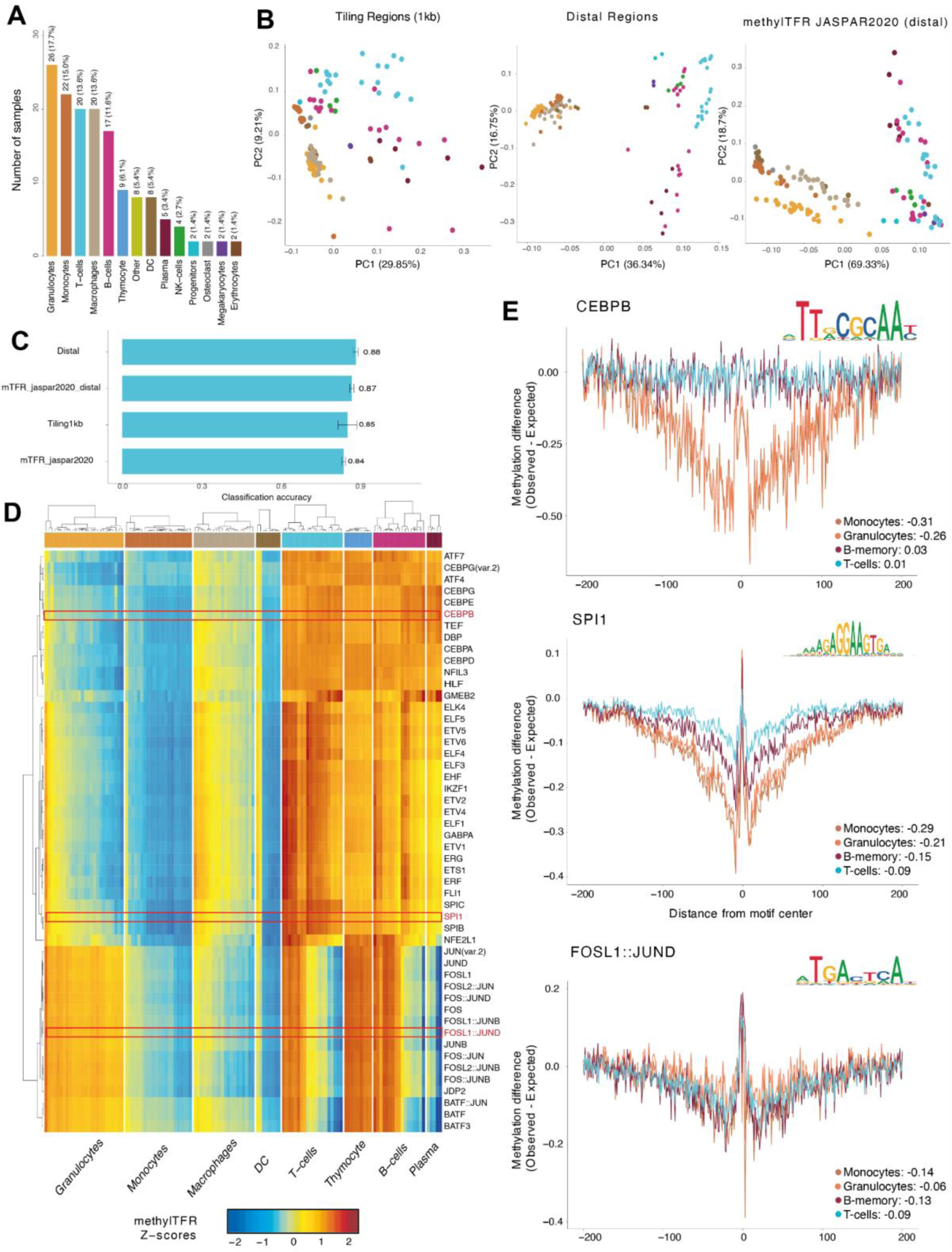
Dissecting the transcription factor dynamics of human immune cell populations A) Overview of cell types and sample numbers in the WGBS dataset from the BLUEPRINT project data release (147 samples across 14 annotated cell types). B) PCA dimensionality reduction of the three representations of the same samples: average methylation in 1-kb tiling regions (left), average methylation in distal regions (middle), and GC-bias-corrected TF deviation scores (JASPAR2020 set with distal subsetting) (right), colored by cell types. C) Random forest classification accuracy for the four representations (genome-wide 1-kb tiling methylation, distal methylation, genome-wide methylTFR, and distal-restricted methylTFR), trained on the first 20 principal components and evaluated over 100 iterations of repeated stratified hold-out validation (20% test fraction). D) Heatmap visualization of bias-corrected deviation scores for the 50 most variable TFs identified by methylTFR. Rows and columns are clustered, columns are annotated by cell type, and three TFs shown in D are boxed in red. E) Bias-corrected motif footprint plots for the three marked TFs (CEBPB, SPI1 and FOSL1::JUND) are shown as the difference between observed and expected methylation around the motif center across cell types, with corresponding methylTFR deviation scores provided in the legend of each panel.

To quantify the predictive power of each representation for cell type, a random forest model was trained on the first 20 principal components of each feature representation and evaluated using 100 iterations of repeated stratified hold-out cross-validation. The mean accuracy was 0.85 for the genome-wide 1-kb tiling representation, 0.84 for genome-wide methylTFR (JASPAR2020), 0.87 for distal-restricted methylTFR, and 0.88 for distal regions (Figure 2C). To confirm the robustness of these findings, we also evaluated the models using a stratified 5-fold cross-validation, which yielded highly consistent accuracy across all representations (Supplementary Figure 1C). This confirms that methylTFR achieves predictive performance comparable to that of a much larger set of distal region features, while relying on a comparatively compact feature set of 632 TF motifs (versus 74,554 distal regions and 2,674,318 genome-wide tiling regions). The motifs that contributed most to this classification, ranked by cell type, recovered the same lineage-defining families (Supplementary Figure 1D). To characterize the behavior of the deviation score across cell types, we examined variable transcription factor programs and the sequence composition term for which they are corrected. A chi-squared test identified 68 TFs with significant variability (adj. p-value < 0.05, variance > 1) (Supplementary Figure 2A, Supplementary Table 1), with the CEBP family member appearing in the first position of the ranking. The expected deviation score, which captures the base composition of the regions a motif falls in, varied around a score of 1 (ranging from 0.995 to 1.020) across the 632 motifs, with FOX and ONECUT motifs at the high end and ZBTB33, EGR1, and KLF9 at the low end (Supplementary Figure 2B). Around half of its variance was constant across samples (51.6%), 45.6% varied by motif and sample, and only 2.8% was contributed by a global offset in that sample’s expected score (Supplementary Figure 2C). Because almost half of the composition term differs between samples, a per-sample correction is required rather than a single global adjustment.

We next investigated the 50 TFs with the highest variance in their corresponding bias-corrected deviation scores (Figure 2D). These variable TF motifs included members of the AP1 family (FOS, FOSL1, JUN, JUND), which showed a relative decrease in deviation scores at TFBS in dendritic cells, monocytes, and macrophages. AP1 motifs showed a bimodal distribution in T and B cells, consistent with these populations comprising both naive and memory cells in B cells and CD4+ and CD8+ cells in T cells (Figure 2D). The CEBP family (CEBPA, CEBPB, CEBPD, CEBPE) and the ETS family (ETS1, ERG, SPI1, ELF1, ETV1) separated myeloid cells from lymphoid cells. These families displayed lower deviation scores at their respective motif loci (active TFs) in monocytes and granulocytes, alongside higher methylation in T and B cells and thymocytes. The association between deviation scores and depletion of methylation at corresponding TFBSs is evident in motif footprint plots, which display aggregate methylation profiles around TFBS occurrences (Figure 2E). Methylation around CEBPB motifs was depleted in monocytes and granulocytes (-0.31 and -0.26) and unchanged in B and T cells (0.03 and 0.01). SPI1 was depleted across all four populations, most strongly in monocytes and granulocytes (-0.29 and -0.21, respectively) and least in T cells (-0.09). FOSL1::JUND was depleted to a similar extent in all four cell types, indicating no separation of the myeloid and lymphoid compartments.

PCA separation of methylTFR deviation scores grouped B-cell clusters by their naive-to-memory state, whereas the separation among T-cell samples reflected CD4+ versus CD8+ identity rather than memory state (Supplementary Figure 2D). We further examined the motif footprints of TFs with variable deviation scores to analyze lineage divergence between T and B cells. SPI1 motifs exhibited higher methylation in T cells than in B cells (T cell deviation score: -0.09, compared to -0.14 and -0.15 in naive and memory B cells, Figure 2E), and the observed profiles departed from the expected ones in the same direction (Supplementary Figure 2E). BATF motif methylation was depleted in memory B cells and T cells (-0.15, -0.10) and less so in naive B cells (-0.04), while PAX1 showed B cell-specific methylation activity (Supplementary Figure 2F).

The JASPAR2020 set contains many highly similar motifs, so a single TF family is represented repetitively. To reduce this redundancy, we repeated our analysis using the Altius motif archetypes [2], which group similar motifs into a single representative motif family. Bias-corrected archetype scores separated the cell types similarly, although with lower variation captured in the first principal components (Supplementary Figure 2G). The ccaat_cebp family showed deviation scores indicating methylation depletion in monocytes and granulocytes (-0.04 and -0.05) compared with B and T cells (-0.01 in both), consistent with the member-specific analysis (Figure 2E). The ap1_1 family showed a similar small decrease in all four cell types, as we observed for FOSL1::JUND (Supplementary Figure 2H, Figure 2E). Collapsing a family into a single representative motif preserves the signal direction and the ordering of cell types but weakens them, since the archetype encompasses the binding sites of the entire family. Archetypes are therefore the better choice for a non-redundant overview of families, whereas the full JASPAR set is better suited to delineating motifs among family members.

### Joint modeling of methylation footprints and regulon-inferred activity identifies lineage-specific regulators in human immune cells

The corrected deviation scores should reflect methylation at binding sites and not the base GC composition of those sites. We evaluated this across 58 BLUEPRINT samples with paired RNA-seq data, averaging the observed and expected deviation scores for each motif across samples and correlating them across motifs. We hypothesized that a factor that is not expressed does not exhibit binding-specific methylation profiles, and that the correlation between its motif deviation score and the expected score calculated from GC base content likely reflects bias. We therefore used the lowest quartile of mean TF expression as a proxy for unexpressed TFs (n=151) and the highest quartile for expressed TFs (n=152). In the distal motif set, the bias correction reduced this correlation for all motifs (r=0.32 to 0.24), for unexpressed TFs (r=0.42 to 0.24), and for expressed TFs (r=0.20 to 0.15) (Supplementary Figure 3A). The correction acted on both expressed and unexpressed TFs, consistent with the removal of a composition-driven bias independent of binding. In the genome-wide set, which includes CpG islands, the uncorrected scores already decreased with the expected score (r = −0.40 and −0.64 after the correction; −0.24 to −0.71 for unexpressed and −0.56 to −0.73 for expressed TFs).

The transcriptional level of a TF is a weak proxy for how much it regulates targeted gene expression [8], since activity also depends on translation, nuclear localization, and cofactor availability [8]. We therefore estimated TF activity using VIPER [18], which infers a factor’s activity from the coordinated expression of its regulon. TF mRNA and VIPER activity exhibit comparable cell-type variance (median R² of 0.41 and 0.44, paired Wilcoxon padj = 0.072; Supplementary Figure 3C) across 156 TFs available in both. However, VIPER-defined activities exhibited higher correlation coefficients with methylTFR deviation Z-scores than TF expression levels alone (median |r| = 0.30 and 0.19, respectively, paired Wilcoxon padj = 7.4e-07) over 199 motifs (Supplementary Figure 3D). As regulon-based activity aggregates the expression of dozens of targets, it is less sensitive to sampling noise of a single, often lowly expressed transcript, and it reflects the post-transcriptional control that determines how much of a factor is available to bind [18].

To obtain a multimodal quantification of TF activity, we performed factor analysis using MOFA [25], integrating methylTFR deviation scores and VIPER activities from paired samples [18]. The five factors most strongly associated with cell-type variation (R² > 0.75) were selected for further evaluation (Supplementary Table 2). MOFA assigns every feature a weight on every factor, which describes how strongly the methylTFR score of a motif or the regulon activity of a TF contributes to that factor. To determine which modalities contribute most to each factor, we used the feature weights and plotted their relative contributions (Figure 3A). methylTFR deviation scores were weighted more heavily than regulon-derived activities across the five factors, which explained the largest portion of the dataset’s variance and contributed at least 70% of the total weight. Factor 8 had the highest methylTFR contribution, with 92.3% of the absolute weight, followed by factor 2 (85.6%). Associating sample factor loadings with annotated cell types revealed which lineages each factor separates (Figure 3B). Factor 2 (R² = 0.94) separated the myeloid compartment, factor 7 (R² = 0.92) separated B-cells, factor 6 (R² = 0.92) separated granulocytes, and factor 10 (R² = 0.77) separated DC cells and megakaryocytes. Factor 8 (R² = 0.77) did not separate the compartments. Furthermore, combining multiple factors separates cell states in multidimensional space (Figure 3C).

**Figure 3:**
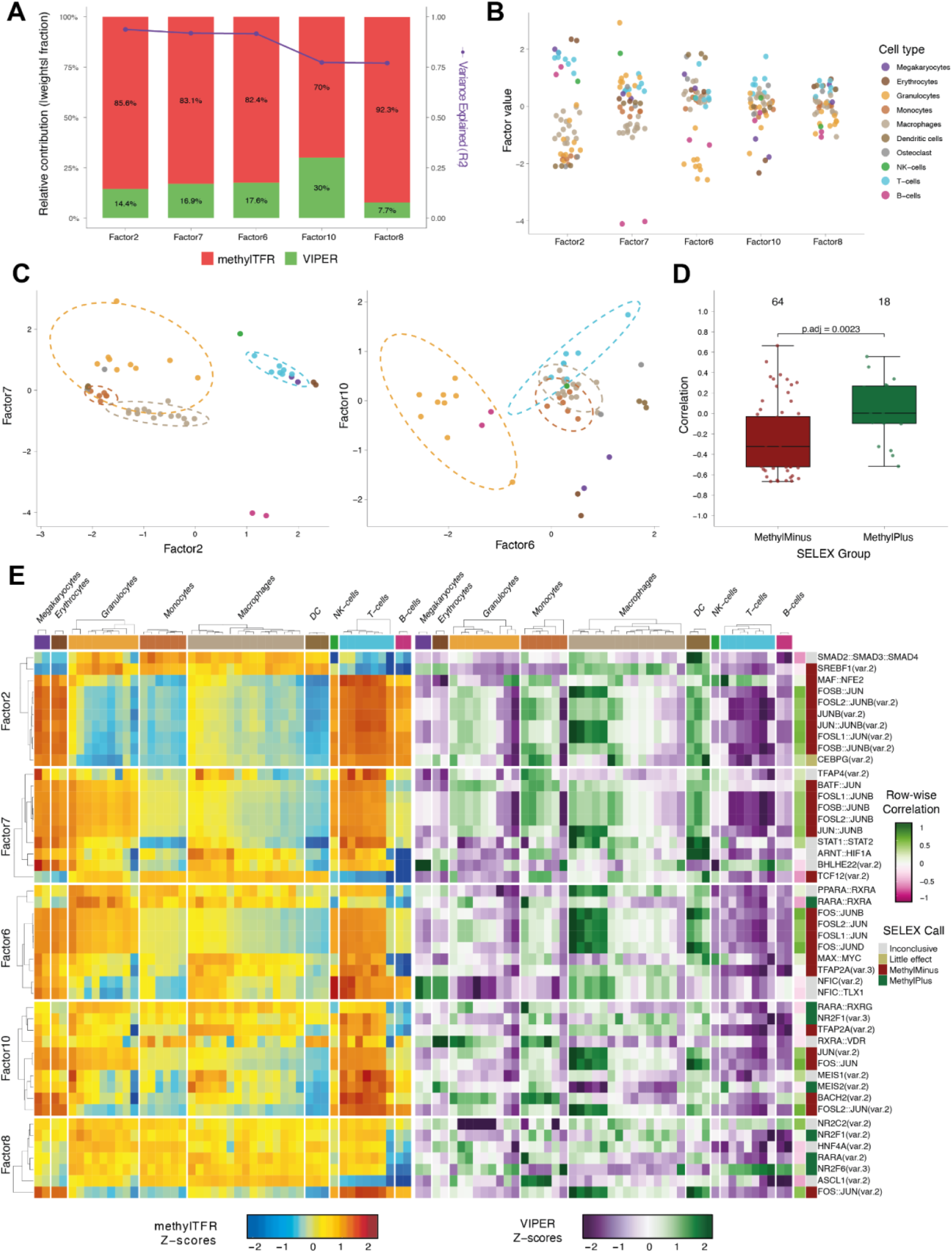
Multimodal factor analysis of binding-site methylation and TF activity across hematopoietic cell types A) Bar plot representing the relative contribution of each modality (red = methylTFR, green = VIPER) for the top five factors most strongly associated with cell type, with variance explained annotated as a purple line. Contributions were calculated using the sum of absolute weights for each modality within the factor. Cell-type variance explained is R² from a one-way ANOVA of factor values against cell type and is distinct from the variance each view reconstructs. Factor 1 was excluded as a global offset. B) Strip plot showing the factor values of five factors with the highest cell-type variance explained. Points are colored by cell type to identify which cell types are associated with which factor. C) Factor-vs-factor plots of factor 2 against factor 7 (left) and factor 6 against factor 10 (right). Points are colored by cell type. Ellipses represent 90% confidence intervals for cell types. D) Boxplot of the Pearson correlation between methylTFR z-scores and VIPER activity across samples, calculated per motif and grouped by the methyl-SELEX call of the motif’s TF. The number of motifs in each category is annotated at the top, along with the p-value from a two-sided Wilcoxon rank-sum test with Benjamini-Hochberg adjustment. E) Heatmap visualization of z-scored activities (methylTFR on the left, VIPER on the right) for the ten highest-loading transcription factors associated with each of the top five latent factors. Row annotations give the Pearson correlation between the two views across samples and the methyl-SELEX call.

Because factor 2 was predominantly methylation-driven and separated the myeloid compartment from lymphoid and erythroid samples, this suggests that DNA methylation at target binding sites, rather than TF regulon activity, is the primary axis of regulatory variation distinguishing these lineages. To test whether TFs’ intrinsic methylation preference explains this coupling, we examined the TFs defining the MOFA factors by their methyl-SELEX classification [11]. methyl-SELEX provides an in vitro measurement of how a factor’s affinity for its motif changes when the CpGs within it are methylated [11]. If the presence of methylation inhibits TF binding, these factors are labeled as methylMinus; in contrast, if the presence of methylation promotes TF binding, they’re labeled as methylPlus. The methylMinus motifs (n=64) showed predominantly negative correlations between the methylTFR and VIPER activity scores across cell types, whereas methylPlus motifs (n=18) were centered on zero (two-sided Wilcoxon rank-sum, p.adj = 0.0023; Figure 3D). TFs negatively impacted by methylation in their respective binding sites therefore gain activity in the cell types with decreased methylation activity at their binding sites (denoted by a negative bias-corrected deviation score), while factors insensitive to DNA methylation show no such coupling. Notably, we observed both positive and negative correlation coefficients between VIPER and methylTFR scores of MOFA-factor-associated TFs (Figure 3D,E). AP-1 dimers are associated with factors 2, 6, and 7, with high methylTFR scores and low VIPER activity in T-cells and the reverse in monocytes and macrophages. Factors 10 and 8 are instead explained by nuclear receptor motifs such as NR2F1, RARA::RXRG, and NR2C2, which carry methylPlus calls more often than the AP-1 factors.

Next, we examined whether the motif matches used to compute the deviation scores lie within active regulatory chromatin, as defined by H3K27ac ChIP-seq signal, across the 35 BLUEPRINT samples with matching methylation and transcriptome data. H3K27ac is a histone modification that marks active promoters and candidate enhancers, and its profiles across the TF motif catalog in our methylTFR annotation exhibited lineage-specific patterns (Supplementary Figure 3F, Methods). Fold enrichment was calculated relative to the mean signal 1.5 to 2 kb away from 500 random positions within the distal regions of the same sample. At the distal binding sites of CEBPB, H3K27ac reached 1.47-fold enrichment in monocytes versus 1.03 in T cells, whereas the random distal positions differed less between the same cell types (1.16 and 1.06), consistent with Figure 2E. Similar trends were observed for other TFs: SPI1 showed 1.39-fold enrichment in monocytes versus 1.14 in T cells; RELA, 1.35 versus 1.19; and JUNB, 1.22 versus 1.06. The motif sites over which the deviation scores are computed therefore exhibit higher H3K27ac signal than random distal positions. For all four factors, this excess was larger in monocytes, where their motifs had lower methylTFR deviation scores and were therefore more hypomethylated.

Across 202 motifs, within the level of individual samples, we did not observe a consistent correlation between the mean H3K27ac fold-change signal and either modality (medians -0.06 and 0.01 for the methylTFR and VIPER activity, respectively) (Supplementary Figure 3E). However, this comparison is indirect, since it is based on fold-change quantifications scaled individually for 35 samples across four cell types, and H3K27ac likely indicates the state of the surrounding chromatin rather than the factor’s occupancy itself.

To test the association of TF motif activity with measured TF occupancy more directly, we computed methylTFR deviation scores from ENCODE WGBS methylomes for samples in which TF occupancy had also been profiled by TF ChIP-seq. For SPI1, CEBPB, ETS1, and RELA, TF ChIP-seq peaks with matched methylome data were available for the primary immune cell population, whereas for the remaining factors, we analyzed the lymphoblastoid line GM12878 as the closest-matching cell type (Supplementary Figure 3G). For the primary cells, we took peaks from ReMap 2022, a uniform reprocessing of public ChIP-seq data that includes ENCODE (SP1 and CEBPB in monocytes, SP1 in B cells), and from UniBind, which defines direct binding sites within ReMap peaks (ETS1 and RELA in CD4+T cells). For each TF with available ChIP-seq data and methylTFR binding site annotation, we split its distal binding sites into those overlapping a peak for the corresponding factor and those that do not, matched the two groups on CpG count and GC content, and computed the deviation score for each group separately (see Methods). Across 28 motif and cell type comparisons from 15 ChIP-seq datasets, ChIP-supported sites had lower deviation scores than matched unsupported sites (median difference in deviation score -0.51) and were therefore more hypomethylated than their composition predicts.

To exclude the possibility that this difference reflects lower methylation at ChIP-supported sites across all cell types, we repeated the comparison against a methylome from an outgroup hematopoietic lineage, where the factor is not expected to bind, using monocytes for the lymphoid sets and T cells for the myeloid sets. The difference between supported and unsupported sites decreased from -0.51 to -0.16 and was larger in the methylome of the bound cell type in 25 of 28 comparisons (sign test p ≈ 3 × 10⁻⁵), so most of the effect is specific to those cells and is not a property of the sites (Supplementary Figure 3H,I). Counting each ChIP-seq dataset once, the difference was larger in the bound cell type for all 15 datasets (median −0.49 versus −0.16, sign test p ≈ 6 × 10⁻⁵). The cell line and the primary cells agreed closely (GM12878: -0.50 vs -0.15, n = 23; primary cells: -0.52 vs -0.16, n = 5).

### DNA methylation-derived TF motif activity identifies transcriptional regulators of memory formation in human CD4+T cells

After demonstrating that methylTFR captures global epigenetic variation across human immune cell types, we investigate whether it can distinguish regulatory differences within T cell subpopulations, specifically in memory formation. We used healthy CD4+T cell populations, including naive (T-naive), central memory (Tcm), and effector memory (Tem), which Durek (2016) [13] previously described as following a consecutive trajectory of memory formation (Figure 4A). To assess methylTFR’s potential for differential TF activity analysis, we compared it with an established workflow for identifying DMRs and performing TF motif enrichment analysis on them. Comparing aggregate CpG methylation using RnBeads [23] across 1-kb tiling regions among T cell populations revealed extensive genome-wide methylation remodeling during memory formation. We identified 1,122 differentially methylated regions (DMRs) between TN and TCM, 1,895 between TN and TEM, and 7,917 DMRs between TCM and TEM (Figure 4B, Supplementary Figure 4A, Supplementary Tables 3 and 4). This asymmetry indicates that most of the substantial epigenetic remodeling along this trajectory occurs during the transition from the naive to the memory state. This pattern is consistent with the progressive, cumulative loss of DNA methylation along the differentiation trajectory described by Durek et al. (2016) [13]. TF motif enrichment analysis of those DMRs using a hypergeometric test implemented in LOLA [24] highlighted TFs associated with activation and memory programs in T cells (Figure 4C, Supplementary Figure 4B). AP1 family factors (JUN, BATF, FOS) were significantly (q-value < 0.05) enriched in memory-associated DMRs that are hypomethylated in memory T cells compared to the naive state. In contrast, SPI and ETS family factors predominated in naive T cells, suggesting a regulatory shift from SPI/ETS-dominated networks toward AP1-driven activity during memory differentiation.

**Figure 4:**
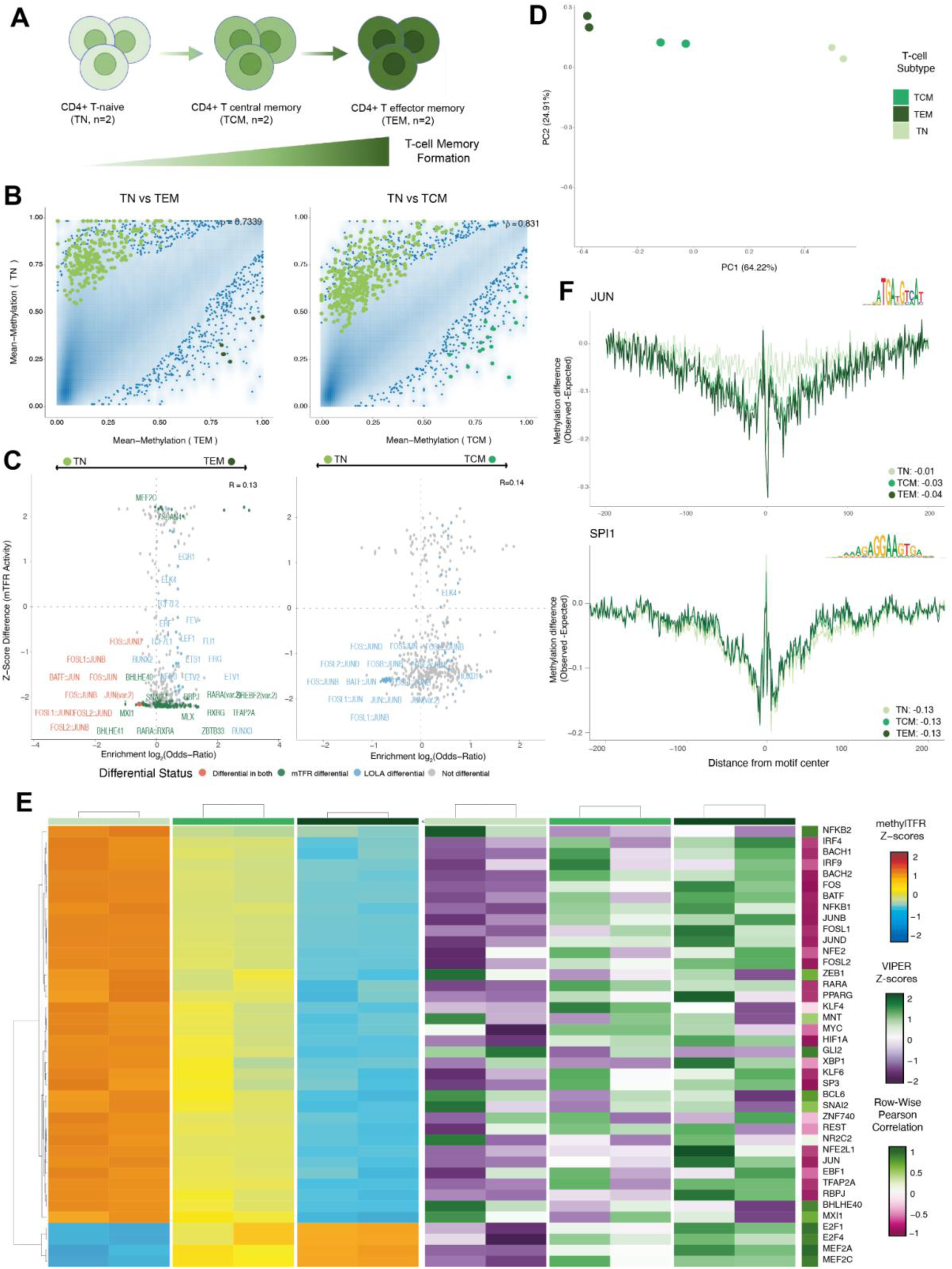
Epigenetic signatures of TF regulation of memory formation in human CD4+T cells A) Schematic differentiation trajectory of memory T-cells. B) Density scatter plots of mean CpG methylation across 1-kb tiling regions comparing the naive state (TN) to effector memory (TEM, left) and central memory (TCM, right) T cells, with differentially methylated regions (DMRs) highlighted. Point density is visualized using 2D kernel density estimation; outliers in low-density regions (<0.1% of all points) are shown individually. C) Differential activity of the JASPAR2020 transcription factor motif set restricted to distal regulatory elements in TN vs TCM (left) and TN vs TEM (right), with significant motifs annotated, shown against the LOLA log2 odds ratios of the same comparison. D) PCA dimensionality reduction based on bias-corrected transcription factor deviations colored by cell types. E) Heatmap visualization of bias-corrected deviation scores for differentially active TFs identified by methylTFR compared with VIPER TF activity of the same TFs inferred using RNA sequencing data. Rows are clustered based on methylTFR deviations. F) methylTFR bias-corrected motif footprint plots of JUN (top) and SPI1 (bottom) motifs in TN, TCM, and TEM cells, annotated with corresponding deviation scores.

While the TFs identified by methylTFŔs differential analysis generally supported the LOLA enrichment results, comparing methylTFR Z-score differences to LOLA log2(Odds-Ratios) revealed that this concordance exhibits notable methodological variance (correlation coefficients: r=0.13 for TN vs TEM and r=0.14 for TN vs TCM; Figure 4C, Supplementary Tables 5 and 6). The two approaches measure different quantities: A hypergeometric test (LOLA) tests whether motif occurrences are overrepresented in DMRs, so a motif contributes only through sites that fall within regions passing the DMR threshold, relative to a selected universe of regions. Most motifs receive odds ratios close to zero. methylTFR summarizes methylation at the center of every distal motif site relative to its flanks, corrected for base composition, and therefore also captures small consistent shifts across thousands of sites outside DMRs. Performing PCA on the deviation scores, the order of the cell types along PC1 (which accounts for 64.22% of the variance) matched the linear progression of memory formation from TN to TCM and TEM (Figure 4D). The methylTFR deviation scores indicated elevated AP1 factor activity in the memory state relative to naive T cells (Figure 4E). We compared methylTFR-derived TF deviation Z-scores with VIPER-inferred regulon activity (VIPER Z-scores) [18] obtained from matched RNA-seq profiles (Figure 4E, Supplementary Table 7). Our findings revealed increased VIPER activity of the AP-1 family (JUN, BATF, FOSL2) in memory subsets (Figure 4E). Although a clear inverse relationship between VIPER and methylTFR deviation scores was evident when comparing naive cells with memory populations, this relationship did not hold for individual memory subsets. The methylTFR activity deviation scores remained comparatively stable between the TCM and TEM states, whereas the corresponding VIPER Z-scores were more heterogeneous, varying both between the two states and between replicates (Figure 4E). This is consistent with DNA methylation acting as a more stable regulatory layer after cell specification, while gene expression varies more between closely related memory states [12,18]. Motif footprint analysis further validated those observations. The AP-1 factor JUN becomes increasingly depleted in methylation across its motif center along the trajectory, from -0.01 TN to -0.03 TCM to -0.04 TEM. The same AP-1 direction was held for FOSL2 and BATF (Supplementary Figures 4C and 4D).

Because LOLA found SPI and ETS motifs enriched in regions hypermethylated in memory cells, thus hypomethylated in naive cells (Figure 4C, Supplementary Figure 4B), we examined the SPI1 motif footprint, which was already depleted of methylation in naive cells and remained unchanged during memory formation (−0.13 in all three states; Figure 4F). Naive cells therefore carry higher methylation across AP-1 motif centers than memory cells, consistent with the findings of Gündüz et al. (2026 [5]. To further examine these findings, we assessed differences in direct TF expression alongside methylTFR and VIPER activity (Supplementary Figure 4E), which highlight the positioning of the representative motifs (JUN, BATF, and FOSL2). Analysis of overall concordance confirmed an expected inverse relationship between methylTFR and both TF expression and VIPER activity at the naive and effector memory stages, weakening to near zero at the central memory stage, whereas the concordance between TF expression and VIPER activity was modestly positive across the same stages, as expected for the two transcriptional measures (Supplementary Figure 4F). Together, our TF activity analysis of memory formation in T cells showed the biological relevance of methylation-based activity estimates and confirmed that methylTFR accurately captures regulatory shifts during T-cell memory formation.

### Integrative multi-modal epigenetic activity analysis reveals key drivers of cell identity from single-cell data

We re-analyzed a publicly available single-cell multi-omics dataset profiling human immune cells exposed to HIV and organophosphate (Supplementary Figure 5A) [5]. Chromatin accessibility-derived (chromVAR) [9] and DNA methylation-derived (methylTFR) TF motif activity broadly matched annotated cell types (Supplementary Figure 5B). Single-cell methylomes cover only a small fraction of CpGs per cell, so most motif sites carry no reads in individual cells. We therefore aggregated cells into pseudobulk samples (metacells). This yielded 266 pseudobulk profiles across 38 samples and seven cell types. The labels used to construct the pseudobulk samples were obtained from the FACS-sorted integration described in Gündüz et al. (2025) [5]. Because the methylation and accessibility pseudobulks were built from the same sample and cell-type labels, each sample was matched across the two modalities, enabling vertical integration.

Using these pseudobulk profiles, we performed MOFA-based integration [25] to identify shared regulatory factors that shape immune cell identity (Supplementary Table 8). The five factors with the highest cell-type R² (0.47 to 0.88) were selected. Factor 3 was most strongly associated with methylation, with methylTFR contributing 96.1% of the absolute feature weights, whereas chromVAR contributed most to factors 1, 2, and 7 (63.3% to 66.8%), and both modalities contributed equally to factor 4 (51.5% methylTFR) (Figure 5A). The identified factors corresponded to distinct immune lineages. Factor 1 (R² = 0.88) separated monocytes, factor 2 (R² = 0.85) separated B cells, factor 3 (R² = 0.57) separated naive from memory T cells, factor 4 (R² = 0.52) separated memory CD8+ T cells from naive T cells, and factor 7 (R² = 0.47) separated NK cells (Figure 5B).

**Figure 5:**
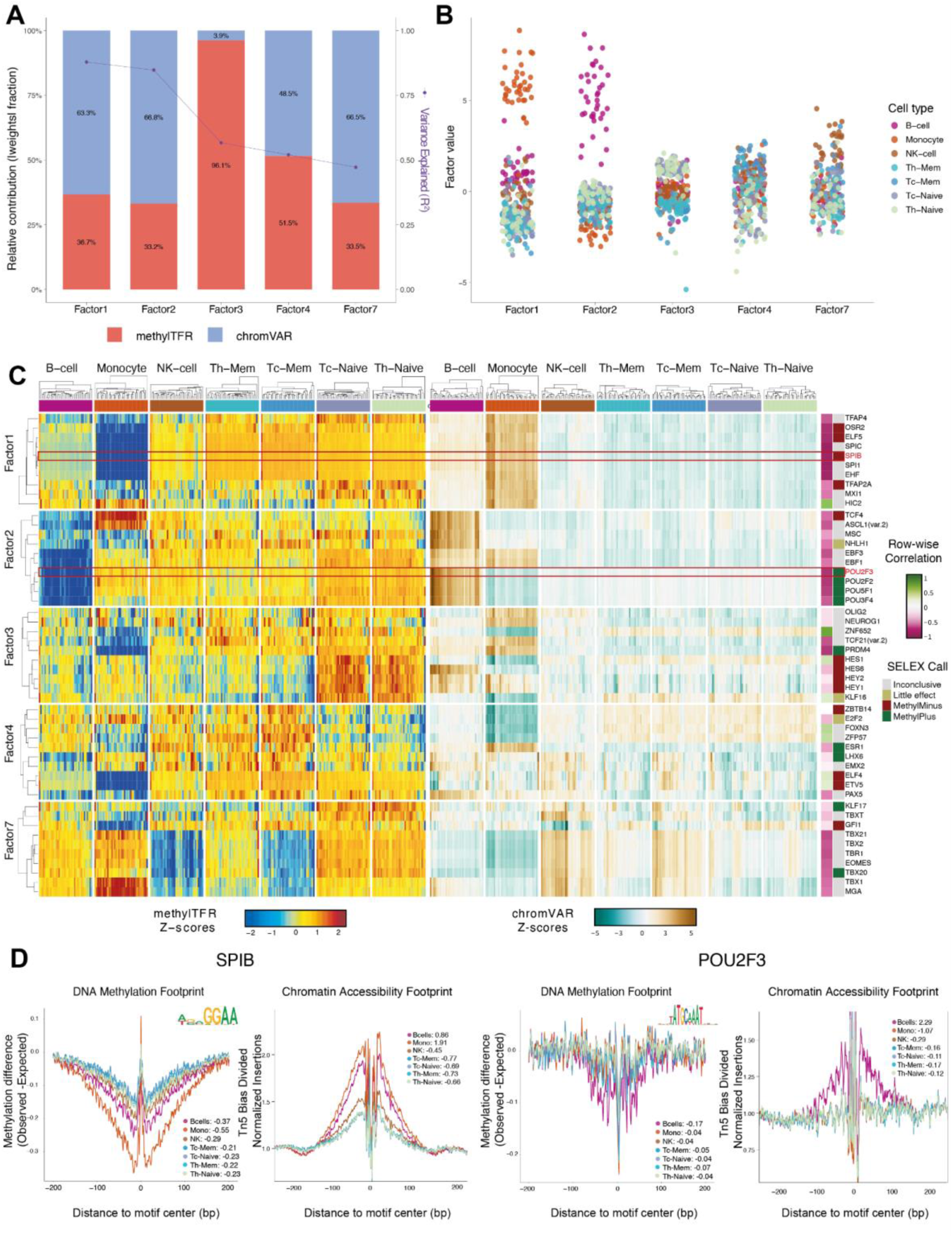
Integrative multi-modal activity analysis in pseudobulked human immune cells A) Bar plot of the relative contribution of each modality (methylTFR and chromVAR) to the top five factors, calculated as the share of summed absolute feature weights, with the cell-type variance explained (R²) overlaid as a line. B) Scatterplot showing the factor values of the five factors with the highest cell-type R² score. Points are colored by cell type to identify which cell types are associated with which factor. C) Heatmap visualization of z-scored activities (methylTFR on the left, chromVAR on the right) for the ten highest-loading transcription factors associated with each of the top five latent factors. D) DNA methylation and chromatin accessibility footprint plots of two example TFs (SPIB left, POU2F3 right) with annotated methylTFR deviation scores.

We further compared the chromVAR and methylTFR activities of factor-associated TF motifs (Figure 5C). Most factor-associated motifs exhibited negative correlation coefficients between methylation- and chromatin-derived TF activity across cell types. In T cells, several TFs had notably high methylTFR Z-scores: key regulators of alternative lineages, such as EBF1, POU2F2, SPI1, and SPIB, exhibited strongly negative chromatin-derived activity scores, consistent with their targeted epigenetic silencing in T cells. For most factors, then, methylation at the TF-binding site was associated with reduced accessibility and potentially with lower expression of candidate target genes. This was also evident when we added methylation-sensitivity labels to the TFs, indicating whether methylation at a TFBS promotes (methylPlus) or inhibits (methylMinus) TF binding [11]. methylMinus TFs outnumbered methylPlus TFs and showed a negative correlation between accessibility and methylation, while methylPlus TFs exhibited a generally more positive correlation (Supplementary Figure 5C). The corresponding motif footprint plots validated these correlations (Figure 5D). A decrease in methylation around the transcription factor binding site leads to increased accessibility at the binding site of the same TF, as was the case for B cells (methylTFR deviation: -0.37, chromVAR deviation: 0.86) and monocytes in SPIB (methylTFR deviation: -0.55, chromVAR deviation: 1.91). Conversely, an increase in methylation leads to reduced accessibility of the same TF, as seen in the T-cell population (e.g., Th-Naive) in the SPIB motif (methylTFR deviation: -0.23, chromVAR deviation: -0.66). The same pattern was observed for the POU2F3 motif. The anticorrelation was especially strong for TFs associated with latent factors 1 and 2, which predominantly consisted of members of the ETS/SPI and POU families, respectively. Both families carry lineage-defining regulators of hematopoietic identity that drive B and monocyte cell lineages, consistent with coordinated demethylation and chromatin opening at their target sites. Averaged across all 629 TFs, the negative correlation was strongest in B cells (median = −0.35), followed by naive T cells and monocytes, and was close to zero across the memory T-cell subsets (Supplementary Figure 5D), where methylTFR detected distinct methylation patterns even when chromatin accessibility remained unchanged. This fits the small accessibility changes we see during memory formation relative to the sharper methylation changes and supports the idea that methylation varies across cell states largely independently of single-cell accessibility, consistent with methylation acting as a more stable, lineage-recording layer than accessibility [12]. Breaking down the correlations by factor and cell type showed that the B-cell factor (factor 2) had the strongest negative correlation in B cells, whereas correlations in memory T cells were near zero or positive across factors (Supplementary Figure 5E).

## Discussion

DNA methylation is a crucial epigenetic modification that regulates transcription factor binding dynamics. However, existing computational approaches for quantifying motif-based TF activity have largely been limited to gene expression or chromatin accessibility data. Here, we present methylTFR, which quantifies transcription factor activity by aggregating DNA methylation signatures around transcription factor binding sites (TFBSs) while accounting for GC bias. methylTFR integrates into established data analysis workflows and is fully compatible with the Bioconductor environment. We compiled motif set annotations for the human genome (Hg38) and Mus musculus (Mm10), which are available as supporting annotation packages (methylTFRAnnotationHg38, methylTFRAnnotationMm10).

While many studies investigate DNA methylation in gene or promoter regions, most candidate regulatory regions that exhibit epigenetic variability across cell types are located in gene-distal regions of the genome [26]. Accordingly, when restricting methylTFR to gene-distal motifs, we observed increased variation attributable to lineage. This indicates that units of distal regulatory elements, rather than individual genes, define cell state. To illustrate the utility of methylTFR for cell state identification and for multi-modal data integration, we applied it to three different use cases: (i) whole-genome bisulfite sequencing (WGBS) data from healthy human immune cells [20], (ii) CD4+T memory cell specification [13], and (iii) metacells derived from single-cell methylomes of human immune cells under HIV and OP exposure [5]. First, we demonstrated that methylTFR deviation scores, despite being based on a relatively low-dimensional yet interpretable set of TF motifs, capture cell-type-specific signatures and perform comparably to or better than larger feature sets, such as distal and tiling regions, in identifying cell type. methylTFR thereby provides a more interpretable overview by reporting aggregated TF activity rather than locus-level change. We also benchmarked the deviation scores against orthogonal measurements of binding, using H3K27ac ChIP-seq at the motif sites and TF ChIP-seq peaks from publicly available data of primary samples and cell lines. H3K27ac, an active-chromatin mark, was elevated at a motif’s binding sites in the lineage where methylTFR calls the factor active. Measured TF occupancy agreed with methylTFR, since a factor’s ChIP-supported sites were more hypomethylated than its sequence-matched unsupported sites.

To further validate the methylTFR algorithm, we identified the TFs with the most variable methylTFR deviation scores in the human immune cell dataset. Our analysis revealed distinct groups of TFs with notably different methylation patterns across cell types, especially between myeloid and lymphoid populations. TF footprints detail where this separation arises: the observed methylation falls below the expected profile in the central bases of the motif and returns to it in the flanks. Two independent assays support the interpretation that these sites are bound by TFs. H3K27ac at the distal binding sites of a motif follows the lineage in which the factor is active, and motif sites overlapping a ChIP-seq peak for the corresponding TF are more hypomethylated than CpG- and GC-matched sites without one. Neither assay measures methylation and binding in the same cells, but both are consistent with the assumption underlying the score: the motif sites in methylTFR reads that are active are the ones the factor binds.

Loyfer et al. profiled 39 human cell types by deep WGBS and applied motif enrichment to the most cell-type-specific unmethylated regions, recovering CEBP, AP-1, and ETS motifs in granulocytes, ETS and RUNX in T-cells, and EBF and E-box motifs in B-cells [27]. A highly similar set of families constituted the most variable motifs in our analysis of human immune cells. Two approaches differ in the unit of analysis. Motif enrichment tests for overrepresentation within a set of regions called unmethylated in one cell type, and therefore return a list of motifs per cell type that depends on the threshold used to define those regions. methylTFR retains motifs as units and returns a continuous score per motif and sample across the entire motif catalog, including factors whose binding sites are neither fully methylated nor fully unmethylated. Such per-sample estimates can be correlated with expression or accessibility for the same factor and used as input to a joint factor model; region-level enrichment supports neither.

We tested the joint effects of DNA methylation and gene expression modules in our paired data from human immune cell populations and identified a latent factor that separated the myeloid compartment from the lymphoid and erythroid samples, with 85.6% of its absolute weight attributable to methylTFR representation. The transcription factors associated with this latent factor belonged to the ETS family, including ETV1, ELF1, ELF3, EHF, and GABPA. Of the two transcriptional representations (TF expression and VIPER), the regulon-based (VIPER) view matched the methylation-derived scores more strongly. Over 199 motifs, VIPER activity correlated with the methylTFR Z-scores at the median absolute r of 0.30, against 0.20 for the mRNA of the factor itself (paired Wilcoxon, padj=7.4e-07), while the two views explained comparable cell-type variance (median R² of 0.45 and 0.42). A single TF transcript is often lowly expressed and noisy, and the amount of protein available to bind is determined post-transcriptionally by translation, localization, and cofactor availability [18]. A regulon aggregates the expression of its targets and therefore reports the consequence of binding rather than the abundance of one transcript, which is the quantity that methylation at the binding sites tracks. Comparisons between methylation-derived activity and gene expression are therefore better made using regulon- or module-level estimates than using raw TF mRNA levels.

In addition to broad cell populations such as T and B cells, TFs associated with differentiation trajectories, such as CD4+ T cell memory formation, can also be characterized using methylTFR. Our results on the differentiation trajectory from TN to TEM cells aligned with those reported by Durek 2016 [13], showing a progressive loss of DNA methylation over time across motif families and a linear differentiation pathway from TN to TCM to TEM cells. VIPER activity was consistent with the observed methylation patterns during memory cell formation: it rose in the AP-1 family and fell in the ETS and SPI families from the naive to the memory state, and the transcripts of the same factors moved in the same direction.

Finally, in our integrative analysis of a single-cell dataset combining accessibility and methylation data, we identified a latent factor (factor 3) to which methylation contributed most, and that was again strongly linked to T cells. This mirrors the pattern we observed in our integrative analysis of DNA methylation and regulon expression: a methylation-dominated factor separates a lineage that the transcriptional view cannot resolve on its own. Integrating accessibility and methylation data revealed that DNA methylation-based TF activity and TF accessibility are complementary, with methylTFR providing additional information on the extent to which changes in accessibility are driven by methylation. Our findings indicate that in human immune cells, changes in accessibility correlate with changes in DNA methylation, and that changes in DNA methylation are manifested more strongly during T cell memory formation than changes in chromatin accessibility.

We also note two limitations of methylTFR: it aggregates the signal across all motif binding sites genome-wide and therefore does not resolve locus-specific activity. That same aggregation is also one of its strengths, since it keeps the output low-dimensional and interpretable, reports regulation at the factor level rather than at the individual locus, and tolerates sparse, unevenly covered methylomes produced by single-cell protocols. Secondly, methylTFR infers TF activity from the methylation state of binding sites rather than directly measuring TF binding. Our integrative analysis showed that the site scores agree with the measured TF occupancy, although binding itself was not observed.

Together, our analysis showed that methylTFR complements the TF expression and accessibility view by providing TF signatures driven by DNA methylation, and we anticipate that it will be a valuable tool for the community to investigate TF activity from DNA methylation data in a wide range of biological contexts. methylTFR, its companion annotation packages, documentation, and use cases are available on the package website (https://epigenomeinformatics.github.io/methylTFR/). It installs as a standard R/Bioconductor package, and its interface for estimating TF activity and downstream analysis integrates with existing methylation analysis workflows.

## Conclusions

Quantifying transcription factor (TF) activity is a valuable tool for assessing changes in gene regulation, yet existing computational tools for DNA methylation analysis have placed limited focus on TF-binding dynamics. Here, we present methylTFR, a method and software package for inferring TF activity directly from DNA methylation signatures at transcription factor binding sites, while correcting for GC composition biases.

Across three distinct use cases (healthy human immune cells, CD4+T cell memory formation, and single immune cell populations under pathogen challenge), we demonstrate methylTFR’s ability to consistently capture biologically meaningful TF activity. methylTFR-derived deviation scores were cell-type-specific, interpretable, and complementary to both chromatin accessibility and gene expression data. Integrative analyses revealed that DNA methylation contributes a distinct, additive layer of regulatory information, particularly in T cell populations, where methylation changes exhibited more pronounced patterns of TF regulatory influence than accessibility during memory cell formation. Taken together, methylTFR bridges a critical methodological gap in the epigenomics toolkit. Framing DNA methylation in terms of TF activity rather than locus-level changes provides a biologically interpretable and scalable framework applicable to bulk and single-cell methylome data. Our use cases illustrate methylTFR’s potential to serve as a valuable resource for the community in investigating TF-driven regulatory mechanisms across diverse biological and disease contexts.

## Methods

### Availability

We implemented methylTFR as an R package under an MIT license to make it publicly accessible and generate reproducible results. The installation instructions and vignette are available in the public GitHub repository (https://github.com/EpigenomeInformatics/methylTFR) and on Zenodo (https://doi.org/10.5281/zenodo.22830103). Package documentation and example use case vignettes are available at https://epigenomeinformatics.github.io/methylTFR/. We provide the code to reproduce our analysis and figures, along with the bias-corrected deviation scores as a methylTFRdeviations object in RDS format at: https://github.com/EpigenomeInformatics/methylTFR_manuscript/. The development version of the methylTFRAnnotationBuilder R package is publicly available under an Artistic-2.0 license at https://github.com/EpigenomeInformatics/methylTFRAnnotationBuilder. The methylTFRAnnotationHg38 package of pre-computed annotations for Hg38 is available at https://github.com/EpigenomeInformatics/methylTFRAnnotationHg38 along with the genome mm10 annotations https://github.com/EpigenomeInformatics/methylTFRAnnotationMm10. The versions of the annotation package methylTFRAnnotationHg38 used in the analysis can be accessed via Zenodo: https://zenodo.org/records/22206980. The methylTFR and accompanying annotation packages are currently under evaluation for Bioconductor development branch 3.24.

### Annotation of transcription factor binding sites

TFBS annotations were derived from the reference motif catalog for the Hg38 genome assembly. The analysis reported here used the JASPAR2020 motif set [3]; the framework also supports Altius motif archetypes [2] and CISbp-v2 [4]. Each TFBS was defined as a symmetric 500 bp window spanning 250 bp upstream and downstream of the motif center.

### methylTFR deviation scores and bias-correction

To quantify deviation scores (e. g., TF activity score) per motif, we computed how DNA methylation at the motif center differs from the local background (Figure 1). The background was estimated from regions located ±250 bp around the motif center, the midpoint of the motif sequence. The deviation score is defined as the mean methylation level in the central ±25 bp of the window (the footprint base) divided by the mean of the two flanking intervals, 200 to 250 bp on either side of the motif center. Because the GC composition of a sequence can bias methylation patterns, we calculated deviation scores from both the observed and expected methylation profiles and used the difference between them as the final bias-corrected value. Here, the observed deviation scores were obtained by aggregating the methylation levels across a sample for all putative TF binding sites (TFBS) and their flanking regions, and the expected deviation scores were calculated analogously for each sample’s expected methylation level for each motif (see next section). Subtracting the expected score from the observed score yields the bias-corrected TF deviation score.

### Expected methylation scores

Genome-wide GC distribution was estimated by dividing the genome (Hg38) into 30-nucleotide windows. For each window, the GC content was defined as the proportion of Guanine (G) and Cytosine (C) nucleotides relative to the window size (e.g., (#G + #C) / window size). Each window was assigned to one of five GC content bins: [0, 0.3), [0.3, 0.36), [0.36, 0.4333), [0.4333, 0.5333), and >0.5333. The resulting genome-wide GC content table was used to compute average methylation levels per sample within each GC bin. In addition, methylTFR requires a GC frequency table of the specified motif set to compute the expected methylation scores of a motif from the samples. To generate the GC frequency table for motif m, each binding site region was expanded to 500 bp plus the motif length and tiled with 30-bp sliding windows, using the reference genome without strand reorientation; windows that overlapped chromosome boundaries were truncated. GC content for each 30-bp TFBS window was calculated as above, and the distribution of these windows across GC bins defined the motif-specific GC frequency table. Expected methylation for motif m was then obtained by multiplying the sample-specific average methylation levels for each of the five GC bins by the motif’s GC-bin frequency table.

### Evaluation of the bias correction

We evaluated the bias correction on 58 BLUEPRINT samples using paired RNA-seq data. For each motif, the observed, expected, and bias-corrected deviation scores were averaged across samples, and the observed score was recovered by adding the expected score back to the stored bias-corrected score. TF expression was computed as log2(CPM+1) from raw RNA-seq counts and averaged across samples. For heterodimer and variant motifs, the highest-expressed subunit was used. Motifs were split by the expression quartiles of the JASPAR2020 distal set: motifs whose TF fell in the lowest quartile as expressed factors. The same cut points were applied to the genome-wide set. For every motif set and expression group, we calculated the Pearson correlation between the expected score and the uncorrected and corrected scores across motifs. To show the trend, motifs were binned into deciles of the expected scores. The uncorrected and corrected scores were each centered on their respective means and averaged per bin, with their standard errors.

### Overview of the methylTFR R package

methylTFR takes two kinds of input: the user’s methylation samples, imported with “read_methylome()”, and the built-in annotation resources methylTFRannotationHg38 or methylTFRannotationMm10 (see Availability), which provide the genome-wide GC distribution, motif GC frequency tables, and transcription factor binding sites (TFBS). Deviation scores are computed for a single TF with “computeDeviation()”, for several samples and motifs at once with “run_methylTFR”, and directly from the RnBeads object using “run_methylTFR_RnBeads()”. All three return a *methylTFRdeviations* object, from which the deviation scores are retrieved with “deviations()” and their row-wise Z-scores with “deviationZScores()”. These are the inputs to the downstream analyses implemented in the package: “computeZScoreVariability()”, which ranks motifs by the variability of their activity, and “differential_deviation_test()”, which identifies motifs with differential activity between groups. Two functions visualize the underlying profiles: “plotExpectedFootprint()” for the observed and expected methylation around the binding sites and “plotMotifFootprint()” for the bias-corrected footprint. Detailed documentation, resources, and case studies are available on the methylTFR GitHub page and the official package website (https://epigenomeinformatics.github.io/methylTFR/index.html).

### Use case analyses

#### Processing DNA methylation samples with RnBeads

We used the RnBeads package (version 2.12.2) to process raw DNA methylation samples and perform quality control [23]. The RnBeads configuration is set with the following parameters: “strand.specific=FALSE”, region.types=c(“distal”, “tiling1kb”), “filtering.sex.chromosomes.removal = TRUE”. The raw data is imported using “rnb.run.import”, followed by quality control and preprocessing via “rnb.run.qc” and “rnb.run.preprocessing”, both with default settings from the RnBeads package. Differential analysis with limma [28] is carried out for all region types using RnBeads’ “rnb.run.differential” function with “differential. comparison.columns” set as “cellTypeShort” and “cell.type” for human immune cells and CD4+T cells, respectively. For memory T cells, LOLA enrichment was performed subsequently using the RnBeads function “performLolaEnrichment.diffMeth” against the LOLA TF Motifs database (Hg38), evaluating differentially methylated regions identified by RnBeads for an overrepresentation of TF binding motifs [24].

#### The bias-corrected deviations with methylTFR

First, the bias-corrected deviation scores were computed using the *“*run_methylTFR_RnBeads*”* function from methylTFR, which takes the preprocessed RnBeads set as data input. The following parameters were used: *“threads = 32”, “chunkSize=15”*, and “ignoreStrand=TRUE”. We applied the same deviation scoring to single-cell methylomes from the ECHO dataset. The input was per-cell allc files rather than an RnBeads set, so deviation scores were computed with the “run_methylTFR” function using “filetype = ‘allc’”, “cov_threshold = 1”, “chunkSize = 10”, “threads = 30”, and “ignoreStrand = TRUE”, against the JASPAR2020 motif set. Only methylation cells overlapping the matched single-cell ATAC-seq cells were retained, and one deviation object was written per cell type. The annotation files were obtained by running the methylTFR functions “getGCfreq(motifSet)” and “getTFbindsites(motifSet)” using the Altius and JASPAR2020 motif catalogs, and “getGenomeGC()”. The resulting file contains the deviation scores for each transcription factor in the motif set. With these scores, we identified transcription factors with significantly different deviation scores between cell types (TN vs TEM/TCM for CD4+T cells) and the most variable TFs for the blueprint dataset. To identify the differential TF activity, the “differential_deviation_test” function from methylTFR was applied to the raw deviation scores using a two-sided, parametric approach (“alternative = *‘two.sided’”*, *“parametric = TRUE”).* This performed a two-sample t-test for pairwise comparisons between groups (naive T cells vs. TCM/TEM). For the most variable TFs, we applied methylTFR’s “computeZScoreVariability” function. Significance was assessed using a chi-squared test. Only the most significant transcription factors were retained, with adjusted p-values < 0.05 and variability> 1. For those transcription factors, we calculated the z-scores (“deviationZScores()” from methylTFR). We plotted the top 50 factors as a heatmap using the “Heatmap” function from the *“ComplexHeatmap”* package (Version 2.22.0) [29]. To generate the motif footprint plots across all cell types, we first extracted the expected and observed methylation values for each motif using the “plotExpectedFootprint” function offered by methylTFR. The bias-corrected scores were computed as the observed-to-expected methylation ratio and normalized against the mean signal of the outer 50 bp flanking regions. These values were then used to visualize the footprint profile around each TF’s motif center.

#### Inferring TF activity from TF regulons

To infer transcription factor activity of CD4+T cells from gene expression data, we used the R package “VIPER” (Version 1.36.0) [18]. “VIPER” infers the TF activity by analyzing the gene expression of its downstream target genes (regulons). These regulons were obtained from the “DoRothEa” database [19,30], but only regulons with high confidence (levels A-C) were included. The normalized enrichment scores per TF were calculated by running the function “viper” with parameters “method = ‘scale’ and “minsize = 4”, applied to log-transformed TPM-normalized gene expression counts. From this data, we created a gene expression matrix with gene short names as rows, samples as columns, and TPM-normalized counts. The results were visualized again with a heatmap. We retained only transcription factors that were in the differentials we obtained from methylTFR and had more than 0 TPMs, resulting in 40 transcription factors for plotting. We also calculated row-wise Pearson correlations between corresponding samples in the methylation and gene expression data and added the scores as a column to the heatmaps. Pairwise concordance between two modalities was quantified as the product of their per-motif Z-scores within a sample; therefore, a positive value means the two measures move in the same direction, whereas a negative value indicates opposite directions. The distribution of these products across motifs is shown per differentiation stage for CD4+ memory formation.

For the BLUEPRINT samples, TF mRNA (log2 CPM) and VIPER activity were compared in the samples with both methylTFR deviation scores and RNA-seq data. Samples annotated as “other” and “thymocyte” were excluded, along with the cell types with fewer than three samples. Each motif was matched to the first subunits that had both an mRNA and a VIPER measurement. For each TF, the cell-type variance explained by each measure was calculated as R² from a one-way ANOVA against cell type. For each motif, we calculated the Pearson correlation across samples between the methylTFR deviation Z-scores and each of the two measures. The two measures were compared using a paired Wilcoxon signed-rank test, with R² for variance and absolute correlation for agreement. The two p-values were adjusted together with the Benjamini-Hochberg procedure.

#### Pseudobulk construction for the single-cell data

Single-cell methylomes were aggregated into pseudobulk profiles per sample and cell type, using the cell-type labels from the FACS-sorted integration of Gündüz et al. [5]. Cells labeled “Other” were excluded. For each combination of sample and cell type, methylated and total read counts in CpG context were pooled across cells at the single-base resolution from per-cell ALLC files. CpGs on chrX, chrY, chrM, and the lambda control (chrL) were excluded. This yielded 266 pseudobulks across 38 samples and seven cell types. methylTFR deviation scores were computed for each pseudobulk against the distal JASPAR2020 motif set. Row-wise Z-scores of these scores and the chromVAR Z-scores of the matching pseudobulks were used as the two views in the MOFA integration.

#### Inferring TF activity from chromatin accessibility and motif footprinting

Single-cell chromatin accessibility was previously processed with ArchR [32] by Gündüz et al. [5], which computes chromVAR-based TF activity [9]. We used the CCA-matched labels from that work and retained only the cells with overlapping single-cell methylomes. Motif positions were obtained for the JASPAR2020 set with “getPositions(project, name = ‘jaspar2020’)”, group coverages were added per mapped cell type with “addGroupCoverages(groupBy = ‘cell_type_meth’)”, and TF footprints were computed over the selected marker motifs with “getFootprints” and drawn with “plotFootprints”, subtracting the Tn5 insertion bias so the profiles are comparable to the methylTFR footprints. Per-cell-type TF activity was summarized from the chromVAR-adjusted Z-score matrix and shown alongside the footprints.

#### Predicting the cell type with a random forest model

We performed multi-class cell-type classification on four data representations: genome-wide and distal-restricted JASPAR2020 motif deviations computed with methylTFR, and 1-kb tiling and enhancer-regulatory (distal) methylation profiles from RnBeads. Across all representations, dimensionality reduction was performed using the “prcomp” function with parameters set to “center=TRUE” and “scale=FALSE”. We extracted the first 20 principal components (PCs) as input features. The classification is performed using the randomForest package [31] with the parameter “ntree=500”. To evaluate model stability, we carried out two cross-validation strategies. First, a repeated stratified hold-out split is performed over 100 iterations with a 20% test fraction, ensuring singleton classes are retained exclusively in the training set. Subsequently, a stratified 5-fold cross-validation was conducted, excluding the cell types with fewer than 2 samples to ensure valid fold partitioning.

#### Multi-omic factor analysis with MOFA

We performed two separate multi-omic integrations using the R package “MOFA2” (Version 1.16.0) [25]. The first model combined methylTFR deviation scores from the blueprint data with protein activity, recalculated using “VIPER” [18] from matched RNA sequencing samples under settings similar to those used for the memory T-cell data, but using log-transformed FPKM gene expression counts. In the second integration, we coupled single-cell DNA methylation with accessibility data. Input data for the first integration were the methylTFR deviation and the TF activity matrix; for the second, pseudobulk z-score matrices from chromVAR and methylTFR. Both matrices of the first integration were z-scored row-wise before training, so that the two views entered the model on a comparable scale. A MOFA object was created using the “create_mofa()” function. We added cell type and sample name as metadata to the MOFA object, prepared it for training with “prepare_mofa”, and trained it with “run_mofa” using the parameter “use_basilisk = FALSE”, running MOFA in a local Python environment via “reticulate”. After training, samples annotated as “Other” or “Thymocytes” were removed. The resulting factors were extracted using the get_factors() function. To find out which factors are associated with which cell types, we performed ANOVA for each factor individually using “aov()”, with value as the dependent variable and cell type as the independent variable. We computed the R² values for each factor as the ratio of the cell-type sum of squares to the total sum of squares, and selected the five factors with the highest cell-type R² values for further analysis. These were plotted as a strip plot of factor values per sample, colored by cell type. For each of those factors, we extracted the feature weights with “get_weights” and expressed the contribution of each modality as its share of the summed absolute weights, which we plotted as stacked bars with the cell-type R² overlaid as a line. For each latent factor, we selected the top 10 transcription factors, plotted 2 heatmaps for each modality, and calculated the row-wise correlation between them. Motifs were selected one factor at a time in turn, ranked by their absolute weight in the methylTFR view, and restricted to motifs that could be matched to a VIPER regulon; each motif and each TF were used only once. Heterodimer motifs such as FOS::JUNB and variant motifs such as JUN(var.2) were matched to a regulon through their first assayed subunit, and the row-wise correlations were compared between the methylMinus and methylPlus groups with a two-sided Wilcoxon rank-sum test, adjusted with the Benjamini-Hochberg procedure.

#### H3K27ac ChIP-seq analysis

We used H3K27ac fold-change signal tracks from BLUEPRINT samples with matching WGBS and RNA-sequencing data. We restricted the analysis to cell types represented by at least five samples and excluded samples annotated as “other” or “thymocyte” cells. For each motif in the JASPAR2020 set, we selected binding sites that overlapped distal regions, subsampled them to at most 2000 sites per motif, and resized each site to a 400 bp window centered on the motif. Signal was read from the bigwig files with the R package “rtracklayer” version 1.62.0, averaged within each window, and then averaged across the windows of a motif to give one value per motif per sample. Because the fold change is scaled per experiment, we Z-scored these values across motifs within each sample so that the between-motif structure, rather than the sample-level global value, was compared. For each motif, we then calculated the Pearson correlation across samples between this signal and the methylTFR deviation z-scores, and between this signal and the VIPER activity of the matched TF, and tested each correlation distribution against zero using a two-sided Wilcoxon signed-rank test, with the Benjamini-Hochberg procedure applied for multiple testing. For the aggregate profiles, we selected CEBPB, JUNB, SPI1, and RELA as example motifs, drew the profiles for monocytes and T cells, took up to 500 distal binding sites, and extracted the signal in 4kb windows centered on the motif, binned into 40 bins of 100 bp. As a control, we drew 500 single positions uniformly at random from within the distal regions, rather than using the midpoints of those regions, which would have centered the control windows on their own peaks. Each sample was divided by its own mean signal across the remaining random windows, so a value of one is the level of unselected distal chromatin.

#### Transcription factor ChIP-seq analysis

TF ChIP-seq peaks were obtained from ReMap 2022 (uniformly reprocessed MACS2 peaks, Hg38) for SPI1 and CEBPB in primary CD14+ monocytes (GSE31621, GSE98367) and for SPI1 in primary B cells (GSE128834); from UniBind (ChIP-eat TFBS, Hg38, release 2022-09-14) for ETS1 in primary CD4+T cells (EXP032565 to EXP032569) and for RELA in primary CD4+ Th1 cells (GSE62482); and from ENCODE (ENCODE4 pipeline, IDR-thresholded peaks, GRCh38) for twelve factors in GM12878. Of the 22 TFs appearing in our results, 11 have no ChIP-seq dataset in any cell type for which a matching methylome exists. Of the remaining 11, seven are covered only in GM12878, and SPI1, CEBPB, ETS1, and RELA are covered in a primary immune population. Datasets with fewer than 5000 peaks were discarded. Where two or more datasets were available for a factor and cell type, peaks were restricted to regions covered in at least two of them, and all peaks were trimmed to ±250 bp around their center. Each peak set was paired with a whole-genome bisulfite methylome of the same cell type from ENCODE (GRCh38 bedMethyl, CpGs with coverage of at least 5): GM12878 (ENCSR890UQO), CD14+ monocyte (ENCSR017BUL), B cell (ENCSR284TCU), and T cell (ENCSR663MXB). The GM12878 methylome and the GM12878 ChIP-seq derive from the same donor (ENCDO000AAK), and the three primary methylomes are from a single donor (ENCDO661BYS), so contrasts among them are donor-controlled. Binding sites were taken from the JASPAR2020 motif set and restricted to the same distal regions used throughout. A site was called ChIP-supported if its center fell within a peak. For each motif with at least 500 distal sites and at least 100 sites in each group, supported sites were matched 1:1 to unsupported sites, exactly on the number of CpGs and to within 0.02 on GC content in a ±250 bp window, sampling without replacement; supported sites with no available pairings were excluded from both groups. Because the per-motif GC frequency table from “getGCfreq(motifSet)” is computed over all of a motif’s sites, it cannot distinguish subsets of those sites. We therefore rebuilt a subset-specific table for each group using methylTFRAnnotationBuilder’s “processMotifs2Matrix” function, restricting the binding sites passed to it to those in that group, so that the expected term reflects the base composition of the sites actually being scored. Deviations were then computed per group using “computeDeviation” (methylTFR) with the genome-wide GC model from “getGenomeGC()”, and the differences between groups were reported. To separate occupancy from chromatin accessibility, the same site sets were rescored with “computeDeviation” against a methylome from a different lineage, using monocytes for the lymphoid sets and T cells for the myeloid sets.

## Abbreviations

5mC: 5-methylcytosine
AP-1: activator protein 1
ATAC-seq: assay for transposase-accessible chromatin using sequencing
cCRE: candidate cis-regulatory element
ChIP-seq: chromatin immunoprecipitation followed by sequencing
CpG: cytosine–phosphate–guanine dinucleotide
DMR: differentially methylated region
GC: guanine–cytosine
LOLA: Locus Overlap Analysis
MOFA: Multi-Omics Factor Analysis
PCA: principal component analysis
RNA-seq: RNA sequencing
SELEX: systematic evolution of ligands by exponential enrichment
TCM: central memory T cell
TEM: effector memory T cell
TEMRA: terminally differentiated effector memory T cell
TF: transcription factor
TFBS: transcription factor binding site
TN: naive T cell
UMAP: uniform manifold approximation and projection
VIPER: Virtual Inference of Protein activity by Enriched Regulon analysis
WGBS: whole-genome bisulfite sequencing.

## Declarations

## Ethics approval and consent to participate

Not applicable.

## Consent for publication

Not applicable.

## Availability of data and materials

The first use case was based on WGBS and RNA-seq data for human immune cell types, which were obtained from the August 2016 release of the BLUEPRINT project (http://blueprint-epigenome.eu/). ChIP-seq data were available from ENCODE (twelve factors in GM12878), ReMap 2022 (GSE31621, GSE98367, and GSE128834), as well as from UniBind (EXP032565 to EXP032569, and GSE62482). Whole-genome bisulfite methylomes were obtained from ENCODE for GM12878 (ENCSR890UQO), CD14+ monocytes (ENCSR017BUL), B cells (ENCSR284TCU), and T cells (ENCSR663MXB). The raw data used in the second use-case analysis are available at the European Genome-Phenome Archive (EGA) under the project accession number EGAS00001001624 for CD4+T memory cell WGBS data. The chromatin accessibility deviation scores reported in the third use-case analysis were computed from the data obtained from the Gene Expression Omnibus (GEO) under accession number GSE306525. The methylation deviation scores reported in the third use-case analysis were computed from the data obtained from the Database of Genotypes and Phenotypes (dbGaP) under accession phs003204.v1.p1 (https://dbgap.ncbi.nlm.nih.gov/beta/study/phs003204.v1.p1/).

## Authors’ contributions

IBG and SKM implemented the algorithm. IBG and RN carried out the analysis. IBG, RN, and FM wrote the manuscript. FM carried out the project’s design, coordination and supervision. All authors read and approved the final manuscript.

## Acknowledgements

We thank Jörn Walter and the members of the Integrative Cellular Biology and Bioinformatics laboratory for constructive comments and feedback.

## Supplemental Information

## Supplementary Figures

**Supplementary Figure 1:**
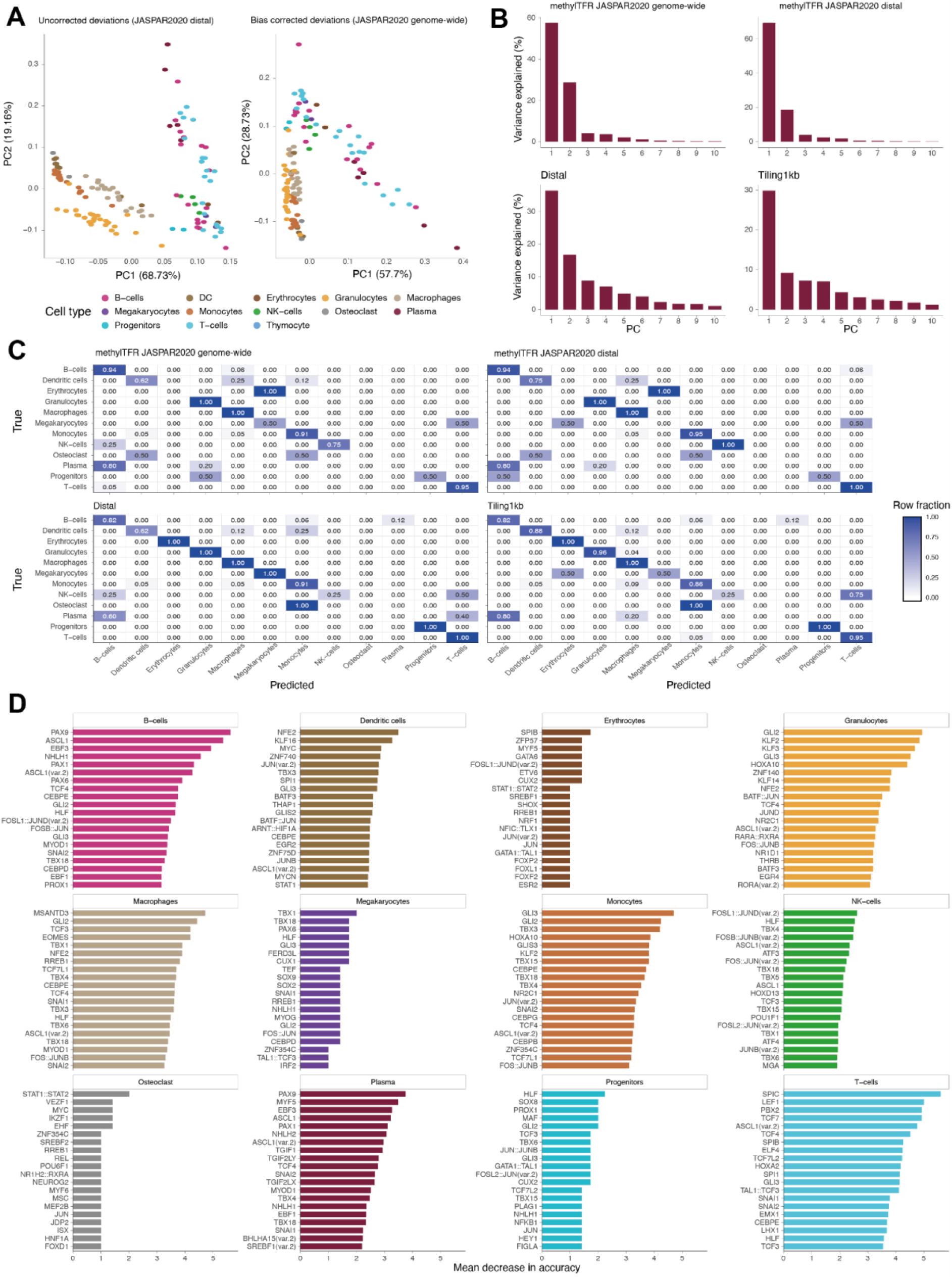
Dimensionality reduction and random forest stability analysis across different methylation representations A) PCA dimensionality reduction of deviation scores computed without the GC bias correction on the distal JASPAR2020 motif set (left) and with the correction on the genome-wide JASPAR2020 set (right), colored by cell types. B) Scree plots of the PCA for the four distinct representations (genome-wide methylTFR, distal methylTFR, distal methylation, and tiling 1kb methylation). Bars give the percentage of variance explained by each of the first ten principal components. C) Confusion matrices showing the classification accuracy of a random forest model trained on the top 20 principal components of each representation using stratified 5-fold cross-validation. Cells are shaded by the row fraction of predicted cell types. D) Top 20 most important motif predictors for each cell type, ranked by their mean decrease in accuracy in a random forest classifier trained directly on the bias-corrected distal methylTFR deviation scores.

**Supplementary Figure 2:**
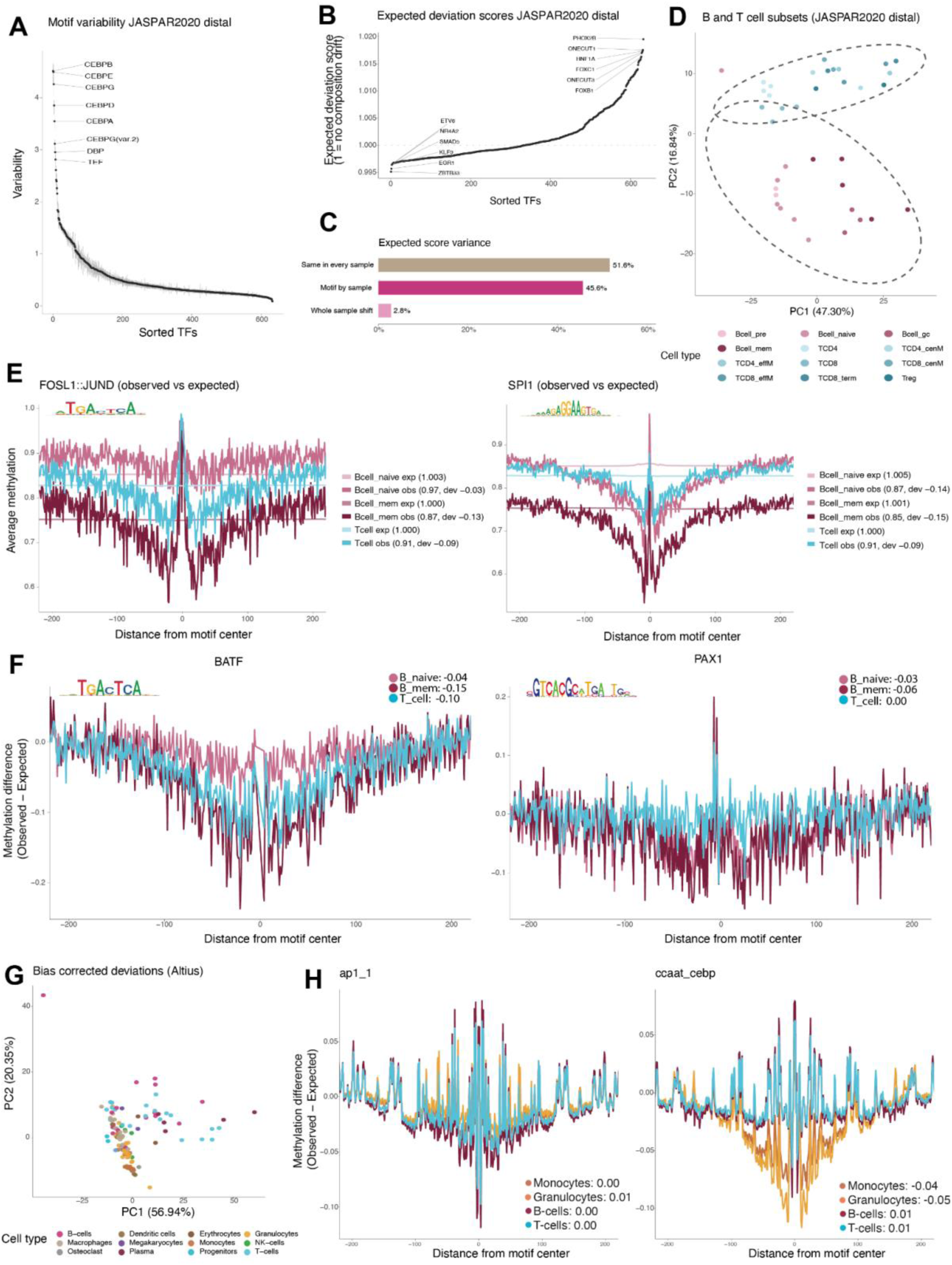
Characteristics of bias-corrected deviation scores and cell-type-specific methylation footprints A) Motif variability of the bias-corrected distal deviation scores, sorted from the most to the least variable TF, with the eight most variable TFs labeled. B) Expected deviation scores of the 632 motifs of the distal JASPAR2020 set, sorted by value, where a score of one indicates no compositional shift. The six motifs at each end are labeled. C) Variance of the expected deviation score, decomposed into three parts: a component that is constant across all samples and depends only on motif composition, a component that varies by motif and sample, and a global offset that shifts all motifs of a sample together. D) PCA dimensionality reduction of the bias-corrected distal deviation scores for the B and T cell subsets colored by individual subgroups. E) Average methylation around the motif center for FOSL1::JUND and SPI1 in naive B-cells, memory B-cells, and T-cells. Observed and expected profiles are shown for each cell type with deviation scores annotated as labels. F) Motif footprints for BATF and PAX1 in the same three cell types, shown as the difference between observed and expected methylation, with the deviation scores in the legend. G) PCA dimensionality reduction of the bias-corrected distal deviation scores computed with the Altius motif sets, colored by cell types. H) Motif footprints for ap1_1 and ccaat_cebp TF families, shown as the difference between observed and expected methylation across monocytes, granulocytes, memory B cells, and T cells.

**Supplementary Figure 3:**
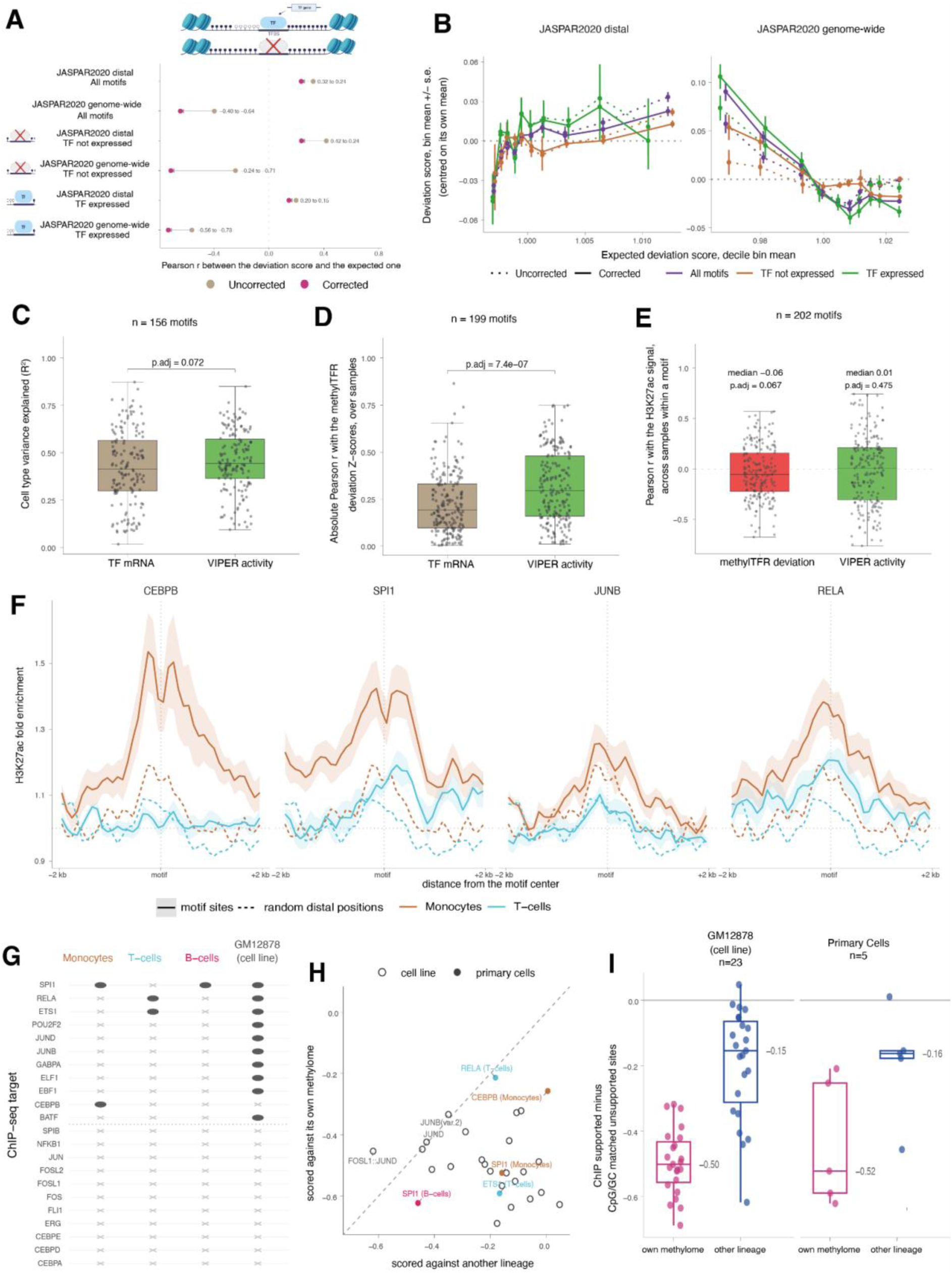
Validation of the corrected deviation scores, the choice of transcriptional view, and the deviation scores against transcription factor occupancy A) Dumbbell plot of the Pearson correlation between the methylTFR deviation score and its expected value, before (tan) and after (magenta) the GC composition correction. Arrows point from the uncorrected to the corrected value. Rows cover two motif sets, distal regions and the genome-wide set of JASPAR2020 motifs, for all motifs, for motifs whose transcription factor falls in the bottom quartile of mean expression, and for those in the top quartile. The dotted line marks zero correlation. B) Deviation scores binned into deciles of the expected score, mean ± s.e. per bin, for the same motif sets and expression groups. Each series is centered on its own mean so that only the slope is compared; dotted lines are uncorrected, and solid lines are corrected. A flat line indicates no residual GC composition dependence. C) Boxplots of the cell-type variance explained (R², by one-way ANOVA of the measure against cell type) by TF mRNA (tan) and by the VIPER activity (green), over 156 TFs carrying both measurements. The annotated p-value is from a paired Wilcoxon signed-rank test, Benjamini-Hochberg-adjusted. D) Boxplots of the absolute Pearson correlation of each measure with the methylTFR deviation z-scores across samples, calculated per motif across 199 motifs. E) Boxplot of the Pearson correlation between the H3K27ac signal at a motif’s distal binding sites and TF motif activity defined by each modality, calculated across samples within a motif, over 202 motifs. Medians and Benjamini-Hochberg-adjusted Wilcoxon signed-rank p-values against zero are annotated above each box. F) Mean H3K27ac profile ± s.e. over samples in 100 bp bins across ±2 kb around the motif center for four example motifs (CEBPB, SPI1, JUNB, and RELA) in monocytes and T cells. Solid lines are the motif’s distal binding sites, and dashed lines are 500 randomly sampled positions within the distal regions, which provide the empirical baseline. Each sample was normalized to its mean signal at 1.5-2 kb from the random positions. Up to 500 binding sites were used per motif. G) Coverage of the occupancy test. Each point represents a transcription factor in a specific cell type in which its binding can be tested. Point area represents the fraction of a motif’s distal binding sites that fall within a ChIP-seq peak, and the fill distinguishes primary immune populations from the lymphoblastoid line GM12878. Crosses mark combinations for which no dataset passes the peak and site thresholds. H) Cell-type specificity of the occupancy effect. For each motif and cell type, the difference in methylTFR deviation between ChIP-supported and matched unsupported distal sites is plotted against the same difference obtained when the identical site sets are scored against a methylome from a lineage in which the factor is not bound. Points below the diagonal are stronger where the factor is bound. I) The two differences from H, shown as boxplots, with the median and interquartile range; each point is one JASPAR2020 motif in one cell type, and the cell line is shown separately from the primary cells.

**Supplementary Figure 4:**
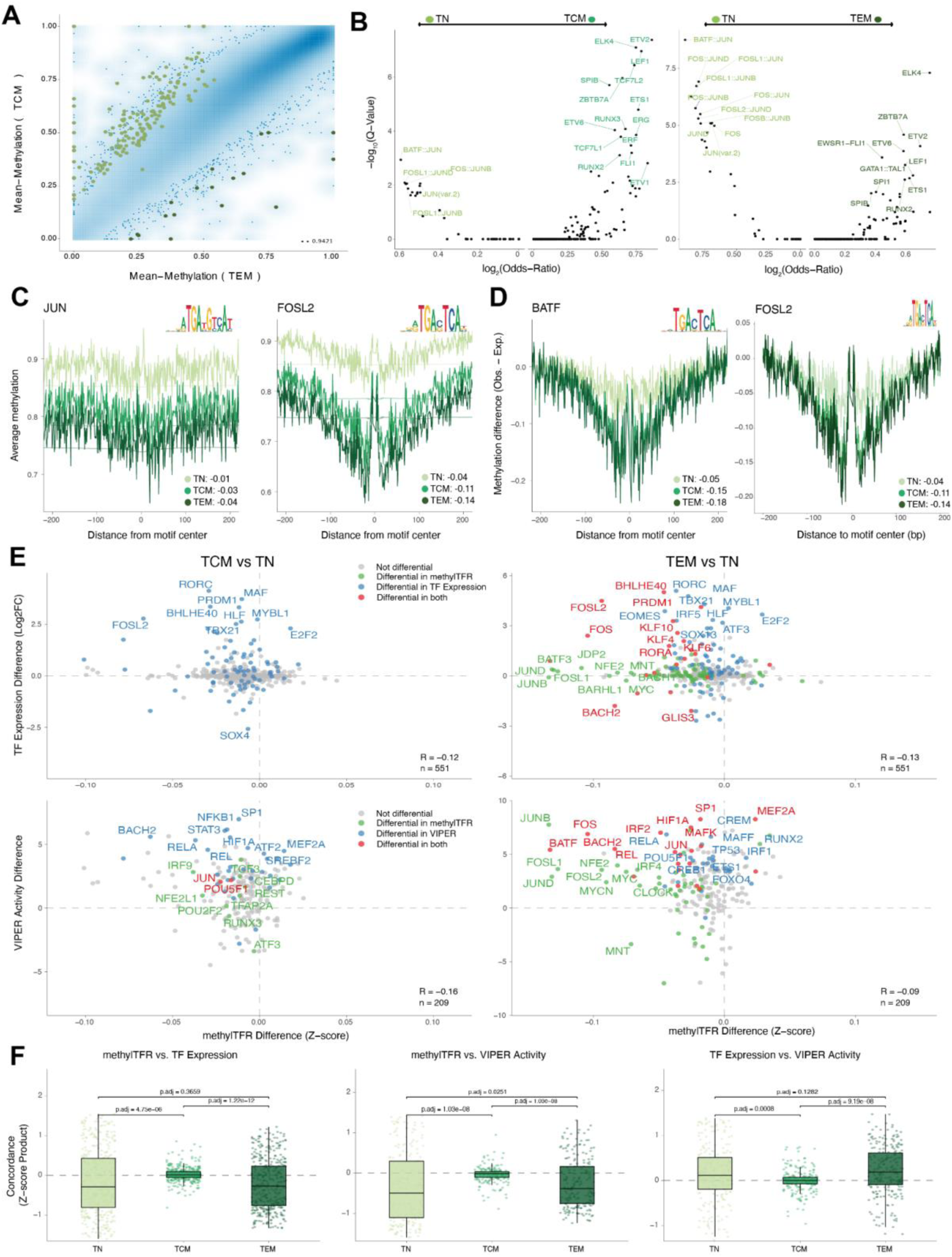
Extended analysis of genome-wide methylation and transcriptional regulatory dynamics during CD4+T cell memory formation A) Density scatter plot of mean CpG methylation across 1-kb tiling regions comparing central memory (TCM) and effector memory (TEM) T-cell populations with differentially methylated regions (DMRs) highlighted. Point density is visualized using 2D kernel density estimation; outliers in low-density regions (<0.1% of all points) are shown individually. B) Volcano plots displaying LOLA TF motif enrichment results within DMRs for the TN vs TCM (left) and TN vs TEM (right) comparisons. C) Motif footprint plots depicting average methylation levels across the motif center for JUN and FOSL2 in TN, TCM, and TEM states. D) Bias-corrected motif footprint plots representing the methylation difference (Observed - Expected) across the motif center for BATF and FOSL2, with observed deviations annotated. E) Scatterplots evaluating the concordance of multi-omics-derived TF activity and expression differences between TCM vs TN (left, TN - TCM) and TEM vs TN (right, TN - TEM). The comparisons shown are TF expression differences versus methylTFR differences (first row) and VIPER activity differences versus methylTFR differences (second row), with the differentially active/expressed motifs highlighted. F) Boxplots quantifying the overall pairwise concordance (calculated as Z-score products) between methylTFR activity, TF expression and VIPER activity estimates across the three differentiation stages.

**Supplementary Figure 5:**
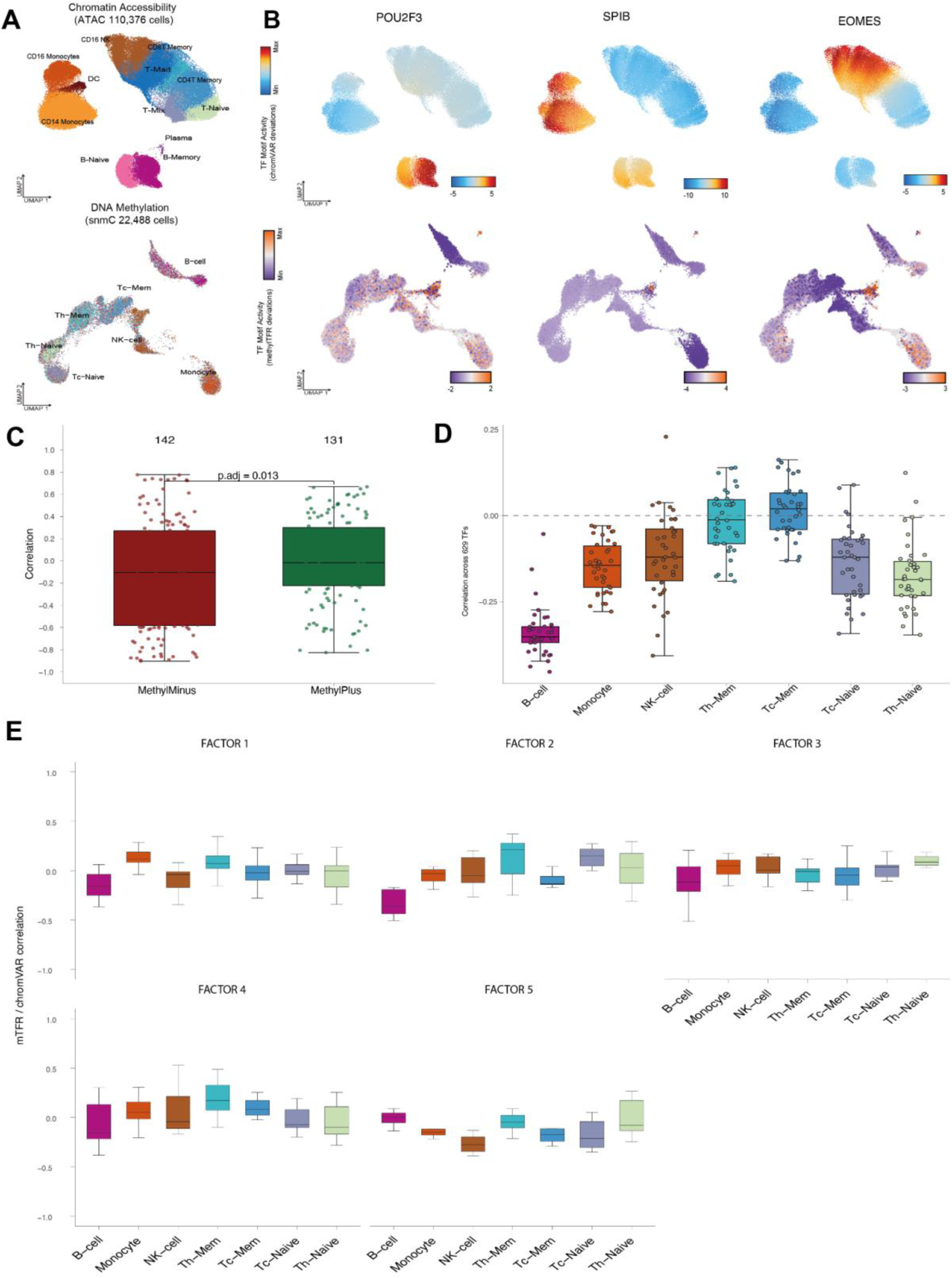
Cross-modal validation of methylTFR against chromVAR in the immune dataset A) UMAP projections of 110,376 cells from single-cell chromatin accessibility (scATAC) colored by cluster labels (top) and 22,488 single-cell methylomes (snmC) colored by cell type labels (bottom). B) UMAP dimensionality reduction colored by the chromVAR TF motif activity (top) and methylTFR TF motif activity (bottom) of POU2F3, SPIB, and EOMES. C) Distribution of methylTFR-chromVAR correlations across TF motifs, grouped by methyl-SELEX call (methylMinus and methylPlus). Each point is one TF, and its value is the Pearson correlation between methylTFR and chromVAR motif activity across pseudobulk samples. Numbers above the boxes give the number of TFs per group; padj is from a two-sided Wilcoxon rank-sum test with Benjamini-Hochberg correction. D) Sample-level agreement between the two modalities, by cell type. Each point represents a pseudobulk sample, and the y-axis shows the Pearson correlation between its methylTFR and chromVAR activity across all 629 TFs. Boxes are colored by cell type, and a dashed line marks zero correlation. E) methylTFR-chromVAR correlations split by latent factor and cell type. For each TF, the correlation is computed within a cell type across its pseudobulk samples: boxes show these per-TF correlations grouped by cell type, with one panel per latent factor. Colors denote the cell type.

## Supplementary Tables

**Supplementary Table 1:** methylTFR transcription factor activity (deviation Z-scores) across human immune cell types.

**Supplementary Table 2:** MOFA feature weights per factor and modality (VIPER regulatory activity and methylTFR deviations).

**Supplementary Table 3:** Differentially methylated distal regions between naive CD4+ T cells (TN) and effector memory T cells (TEM), identified by RnBeads.

**Supplementary Table 4:** Differentially methylated distal regions between naive CD4+ T cells (TN) and central memory T cells (TCM), identified by RnBeads.

**Supplementary Table 5:** Differentially active transcription factors between naive CD4+ T cells (TN) and the TEM and TCM subsets, identified by methylTFR.

**Supplementary Table 6:** LOLA motif enrichment statistics for the pairwise comparisons among TN, TCM, and TEM.

**Supplementary Table 7:** JASPAR2020 distal transcription factors with their methylTFR activity scores, differential status, and matched VIPER activity scores.

**Supplementary Table 8:** MOFA feature weights per factor and modality (chromVAR deviations and methylTFR deviations).

## References

1. Spitz F, Furlong EEM. Transcription factors: from enhancer binding to developmental control. Nat Rev Genet. 2012;13:613–26. 10.1038/nrg3207

2. Vierstra J, Lazar J, Sandstrom R, et al (2020) Global reference mapping of human transcription factor footprints. Nature 583:729–736

3. Fornes O, Castro-Mondragon JA, Khan A, et al (2019) JASPAR 2020: update of the open-access transcription factor binding profiles database. Nucleic Acids Res 48:D87–D92

4. Weirauch MT, Yang A, Albu M, Cote AG, Montenegro-Montero A, Drewe P, et al. Determination and Inference of Eukaryotic Transcription Factor Sequence Specificity. Cell. 2014;158:1431–43. 10.1016/j.cell.2014.08.009

5. Gündüz IB, Wei B, Chen DC, Wang W, Hariharan M, Norell T, et al. Dissecting the epigenome dynamics in human immune cells upon viral and chemical exposure by multimodal single-cell profiling. 2025; 10.1101/2025.09.09.675101

6. Marei HE. Epigenetic regulators in cancer therapy and progression. npj Precis Oncol. 2025;9:206. 10.1038/s41698-025-01003-7

7. Zhang L, Silva TC, Young JI, Gomez L, Schmidt MA, Hamilton-Nelson KL, et al. Epigenome-wide meta-analysis of DNA methylation differences in the prefrontal cortex implicates the immune processes in Alzheimer’s disease. Nat Commun. 2020;11:6114. 10.1038/s41467-020-19791-w

8. Lambert SA, Jolma A, Campitelli LF, Das PK, Yin Y, Albu M, Chen X, Taipale J, Hughes TR, Weirauch MT (2018). The Human Transcription Factors. Cell 172:650–665

9. Schep AN, Wu B, Buenrostro JD, Greenleaf WJ (2017) chromVAR: inferring transcription-factor-associated accessibility from single-cell epigenomic data. Nat Methods 14:975–978

10. Calviello AK, Hirsekorn A, Wurmus R, Yusuf D, Ohler U (2019) Reproducible inference of transcription factor footprints in ATAC-seq and DNase-seq datasets using protocol-specific bias modeling. Genome Biol 20:42

11. Yin Y, Morgunova E, Jolma A, et al (2017). Impact of cytosine methylation on DNA-binding specificities of human transcription factors. Science. 10.1126/science.aaj2239

12. Kim M, Costello J (2017) DNA methylation: an epigenetic mark of cellular memory. Exp Mol Med 49:e322–e322

13. Durek P, Nordström K, Gasparoni G, et al (2016) Epigenomic Profiling of Human CD4+ T Cells Supports a Linear Differentiation Model and Highlights Molecular Regulators of Memory Development. Immunity 45:1148–1161

14. Liu F, Wang Y, Gu H, Wang X (2023) Technologies and applications of single-cell DNA methylation sequencing. Theranostics 13:2439–2454

15. Moore LD, Le T, Fan G (2013) DNA Methylation and Its Basic Function. Neuropsychopharmacology 38:23–38

16. Medvedeva YA, Khamis AM, Kulakovskiy IV, et al (2014). Effects of cytosine methylation on transcription factor binding sites. BMC Genom 15:119

17. Frias AB, Boi SK, Lan X, Youngblood B. Epigenetic regulation of T cell adaptive immunity. Immunol Rev. 2021;300:9–21. 10.1111/imr.12943

18. Alvarez MJ, Shen Y, Giorgi FM, Lachmann A, Ding BB, Ye BH, et al. Functional characterization of somatic mutations in cancer using network-based inference of protein activity. Nat Genet. 2016;48:838–47. 10.1038/ng.3593

19. Gentleman RC, Carey VJ, Bates DM, Bolstad B, Dettling M, Dudoit S, et al. Bioconductor: open software development for computational biology and bioinformatics. Genome Biol. 2004;5:R80. 10.1186/gb-2004-5-10-r80

20. Stunnenberg HG, Consortium TIHE, Abrignani S, et al. (2016) The International Human Epigenome Consortium: A Blueprint for Scientific Collaboration and Discovery. Cell 167:1145–1149

21. Zerbino DR, Wilder SP, Johnson N, Juettemann T, Flicek PR (2015) The Ensembl Regulatory Build. Genome Biol 16:56

22. Kribelbauer JF, Laptenko O, Chen S, Martini GD, Freed-Pastor WA, Prives C, et al. Quantitative Analysis of the DNA Methylation Sensitivity of Transcription Factor Complexes. Cell Rep. 2017;19:2383–95. 10.1016/j.celrep.2017.05.069

23. Müller F, Scherer M, Assenov Y, Lutsik P, Walter J, Lengauer T, Bock C (2019) RnBeads 2.0: comprehensive analysis of DNA methylation data. Genome Biol 20:55

24. Sheffield NC, Bock, C (2016) LOLA: enrichment analysis for genomic region sets and regulatory elements in R and Bioconductor. Bioinformatics 32:587–589

25. Argelaguet R, Arnol D, Bredikhin D, Deloro Y, Velten B, Marioni JC, et al. MOFA+: a statistical framework for comprehensive integration of multi-modal single-cell data. Genome Biol. 2020;21:111. 10.1186/s13059-020-02015-1

26. Ziller MJ, Gu H, Müller F, Donaghey J, Tsai LT-Y, Kohlbacher O, et al. Charting a dynamic DNA methylation landscape of the human genome. Nature. 2013;500:477–81. 10.1038/nature12433

27. Loyfer N, Magenheim J, Peretz A, Cann G, Bredno J, Klochendler A, et al. A DNA methylation atlas of normal human cell types. Nature. 2023;613:355–64. 10.1038/s41586-022-05580-6

28. Ritchie ME, Phipson B, Wu D, Hu Y, Law CW, Shi W, et al. limma powers differential expression analyses for RNA-sequencing and microarray studies. Nucleic Acids Res. 2015;43:e47–e47. 10.1093/nar/gkv007

29. Gu Z, Eils R, Schlesner M. Complex heatmaps reveal patterns and correlations in multidimensional genomic data. Bioinformatics. 2016;32:2847–9. 10.1093/bioinformatics/btw313

30. Garcia-Alonso L, Holland CH, Ibrahim MM, Turei D, Saez-Rodriguez J. Benchmark and integration of resources for the estimation of human transcription factor activities. Genome Res. 2019;29:1363–75. 10.1101/gr.240663.118

31. Liaw A, Wiener M (2002). “Classification and Regression by randomForest.” R News, 2(3), 18–22. https://CRAN.R-project.org/doc/Rnews/.

32. Granja JM, Corces MR, Pierce SE, et al. ArchR is a scalable software package for integrative single-cell chromatin accessibility analysis. Nature Genetics. 2021;53:403–411. doi:10.1038/s41588-021-00790-6

